# A machine learning and network framework to discover new indications for small molecules

**DOI:** 10.1101/748244

**Authors:** Coryandar Gilvary, Jamal Elkhader, Neel Madhukar, Claire Henchcliffe, Marcus D. Goncalves, Olivier Elemento

## Abstract

Drug repurposing, identifying novel indications for drugs, bypasses common drug development pitfalls to ultimately deliver therapies to patients faster. However, most repurposing discoveries have been led by anecdotal observations (e.g. Viagra) or experimental-based repurposing screens, which are costly, time-consuming, and imprecise. Recently, more systematic computational approaches have been proposed, however these rely on utilizing the information from the diseases a drug is already approved to treat. This inherently limits the algorithms, making them unusable for investigational molecules. Here, we present a computational approach to drug repurposing, CATNIP, that requires only biological and chemical information of a molecule. CATNIP is trained with 2,576 diverse small molecules and uses 16 different drug similarity features, such as structural, target, or pathway based similarity. This model obtains significant predictive power (AUC = 0.841). Using our model, we created a repurposing network to identify broad scale repurposing opportunities between drug types. By exploiting this network, we identified literature-supported repurposing candidates, such as the use of systemic hormonal preparations for the treatment of respiratory illnesses. Furthermore, we demonstrated that we can use our approach to identify novel uses for defined drug classes. We found that adrenergic uptake inhibitors, specifically amitriptyline and trimipramine, could be potential therapies for Parkinson’s disease. Additionally, using CATNIP, we predicted the kinase inhibitor, vandetanib, as a possible treatment for Type 2 Diabetes. Overall, this systematic approach to drug repurposing lays the groundwork to streamline future drug development efforts.

## Introduction

With over $800 million spent bringing a single drug to market over the course of 15 years, drug development has remained a costly and time-consuming affair^1^. In response, there has been an increase in interest in drug repurposing, the identification of novel indications for known, safe drugs. Successes in this area have been seen in the past, most notably in sildenafil (e.g. Viagra), which was originally intended to treat hypertension and angina pectoris but was later repurposed to treat erectile dysfunction. Other examples of compounds repurposed for new therapeutic applications include minoxidil^2^ and raloxifene^3^, which are now used to treat androgenic alopecia and osteoporosis, respectively. However, most of these repurposing opportunities were discovered through inefficient approaches including anecdotal observations or hypothesis-driven investigations, and a more efficient approach could lead to many more repurposing opportunities.

Computational approaches for repurposing drugs are appealing in that they can be systematically and quickly applied to many drugs at a low cost compared to their experimental counterparts. One computational approach that has proven to be invaluable in other areas of the drug development pipeline is machine learning. Machine learning is the use of computational algorithms to learn from available data to make novel predictions and gain new insight. Using this technique, one can create unbiased algorithms to match seemingly disparate drugs by comparing their common features^4^, such as clinical indication, toxicity profile^5^ or therapeutic target^6,7^. Previously, our lab used a ‘similarity’ approach, leveraging the principle that similar drugs tend to have similar characteristics, to predict a drug’s target by investigating the known targets of other drugs that were predicted to be “similar” to the investigated drug based on shared features^6^. We found that DRD2, a dopamine receptor, was the predicted target for the compound ONC201. After identifying and experimentally validating this target, clinical trials were shifted to focus on gliomas, which are now successfully completing phase two trials at the time of this publication^8^. The approach of leveraging drug similarity could immensely aid drug repurposing efforts with the appropriate data.

Others have successfully used this ‘similarity’ approach to repurpose drugs and demonstrated high predictive power when tested against FDA approved drug-diseases^9^. However, these methods have primarily linked drugs together using a disease-centric approach instead of using features related to the drug itself (i.e. drug-centric). These repurposing opportunities are identified by predicting diseases similar to the diseases a drug is already known to treat. Disease similarities can be based on semantic, pathophysiological, or clinical similarities related to the drug’s clinical indication. For example, PREDICT, a repurposing method developed by Gottlieb et al.^10^, exploits the semantic similarity of disease terms as a form of disease-disease similarity. Such approaches, while reliable, limit the scope of the repositioning effort in several ways. First, the vast majority of small molecules never reach clinical approval and would be overlooked in this type of analysis. Second, the use of a disease-centric approach biases repurposing predictions toward exclusively similar clinical diseases (i.e.: cancer drugs to other cancer types) ^11^. We postulated that using solely drug information, such as chemical and biological features, would be a more effective and broader approach to drug repurposing.

Here, we propose a novel approach to drug repurposing, which operates by a platform we call, **Creating A Translational Network for Indication Prediction** (CATNIP). CATNIP is a machine-learning algorithm that learns to predict whether two molecules share an indication based solely on the drug’s chemical and biological features, using 2,576 unique drugs. The systematic application of CATNIP to molecule pairs creates a network with ~4.6 million nodes that can then be used to identify potential drug repurposing opportunities. Because CATNIP uses chemical structure and targets as key features, it can effectively bridge between different therapeutic indications. In this report, we have identified various candidate drug classes that are predicted to have therapeutic activity outside of their intended indication in diseases such as Parkinson’s disease and Type 2 Diabetes.

## Results

### Variance in drug indication nomenclature can be standardized

We collected a wide variety of drugs (N=3,066, including both approved and investigational molecules) with a diverse set of indications to ensure that our drug network covered a large portion of the known chemical space. A subset of these drugs (2,576 FDA approved drugs and 2,492 indications taken from DrugBank^12^) were used as a gold-standard of drug-indication associations in the training set for the model. Disease names are often not standardized, which can lead to many diverse names for the same disease. This problem leads to many drug pairs appearing to not have shared indications, when they are associated with two different names for the same disease. To address inconsistencies in nomenclature for drug indications, such as “prostate carcinoma” and “carcinoma of the prostate”, the MetaMap tool^13^ was applied (**Methods**). Using MetaMap, we clustered the 2,492 DrugBank indications into 1,042 standardized indications. A multitude of indication types were included in this standardization including, but not limited to, oncological, mental health, and neurological diseases (**Figure S1A**). Our rigorous standardization of drug indications ensured an accurate training set, allowing for the discovery and modeling of drug-indication relationships.

### Drug pairs sharing indications have other similar characteristics

We hypothesized that pairs of drugs that shared at least one indication would have other similar drug characteristics (**Table S1**). To test this hypothesis, we integrated the similarity of two drugs across chemical and biological drug properties, and created a computational model to predict if two drugs will share an indication (**Figure 1**). All 16 of the drug similarity features (**Table S1**) collected could significantly distinguish between drug pairs known to share an indication and those not known to share an indication (**Figure S2-5**). For example, we found that drug pairs with a shared clinical indication, according to their listed DrugBank indications, tended to have significant overlap in targets (D-statistic = 0.168, p-value < 0.001, **Figure S2A**). The feature which best discriminated between drug pairs that shared a clinical indication versus drug pairs that do not was the similarity between the KEGG pathways that each drug’s targets are involved in (D-statistic = 0.241, p < 0.001, **Figure S4C**). Pathway similarity was calculated as the Jaccard Index between the KEGG pathways that contain each drug’s gene targets (**Methods**). The difference in effect size between the target similarity and the pathway similarity (D-statistic= 0.168 vs 0.241, respectively) indicates that the drugs do not necessarily have to target the same exact genes, but rather the same biological pathway, in order to share a clinical indication. Additionally, we found that drug pairs that share an indication had a more similar chemical structure than drug pairs that did not share an indication (D-statistic = 0.105, p-value < 0.001, **Figure S5A**). Overall, these features seem to indicate sufficient power in differentiating drugs that share and do not share indications, which we hypothesized can then be leveraged to create a predictive model.

**Figure 1:**
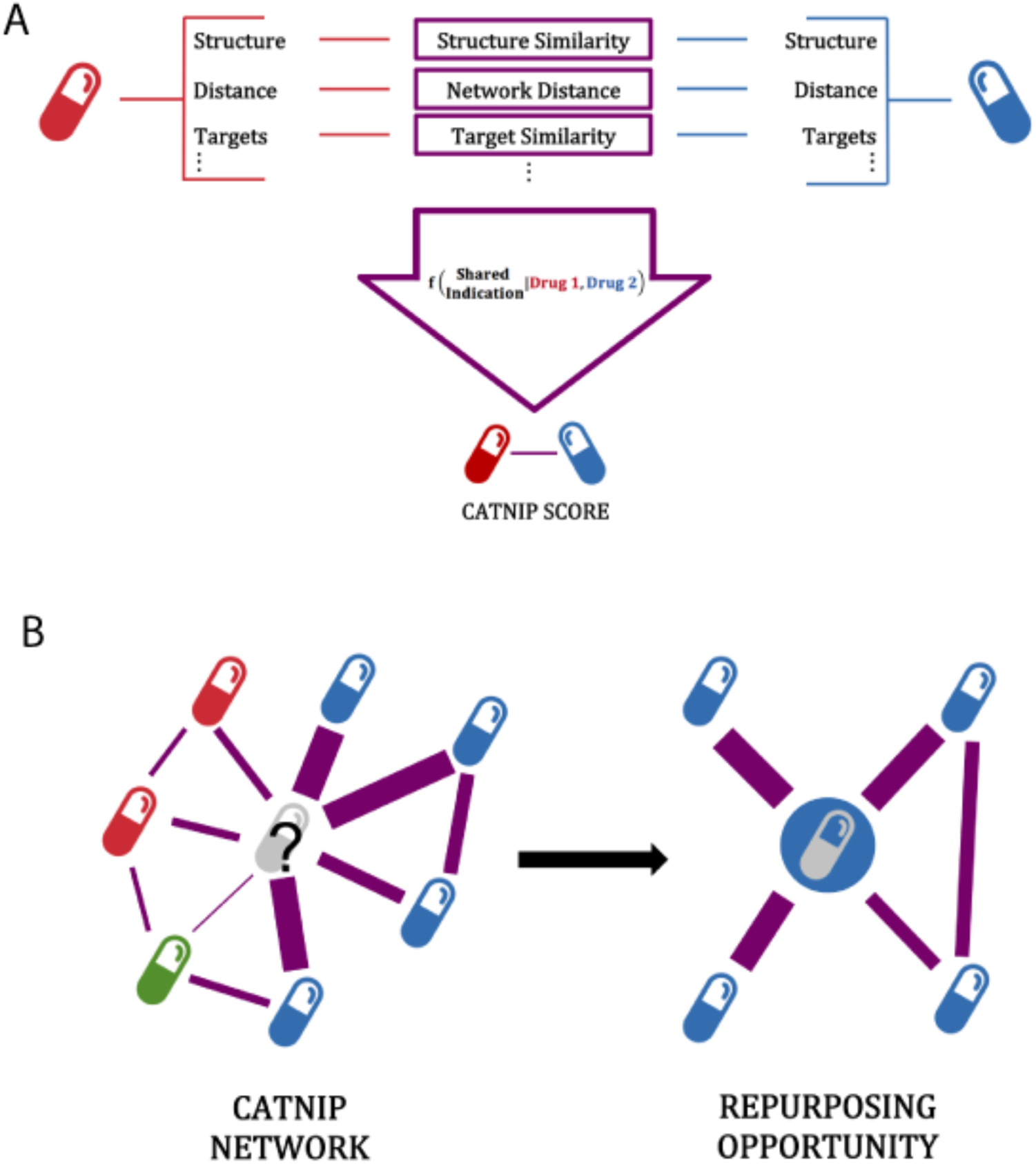
Schematic of CATNIP repurposing approach. A) The use of drug similarity properties to predict if two drugs will share an indication using a gradient boosting model, the model is referred to as CATNIP. B) Schematic showing the use of CATNIP output scores to create a network, with the scores used as edge weights. The colors of each drug represent the known disease and this demonstrates how one could identify novel indications for drugs through the network.

### Drug pairs that share indications can be predicted by model

Using these diverse drug properties as features we trained a Gradient Boosting model to predict if two drugs share a clinical indication. A Gradient Boosting model showed superior results when compared with other algorithms (**Methods, Table S2**). The model output is a drug similarity score (hereby referred to as a “CATNIP score”), which allows us to classify drug pairs that share clinical indications. We performed a 5-fold cross-validation analysis and achieved significant predictive performance with an area-under-the-receiver-operator curve (AUC) of 0.841 (**Figure 2A**). We confirmed the statistical significance of our model with a precision-recall curve (PRC) because of the class imbalance in our dataset between drug pairs that share indications against those that do not (23,840 Shared, 1,299,623 Not Shared). When compared to random predictions, our model showed significant improvement (0.189 vs 0.0184 area-under PRC, **Figure S6**).

**Figure 2:**
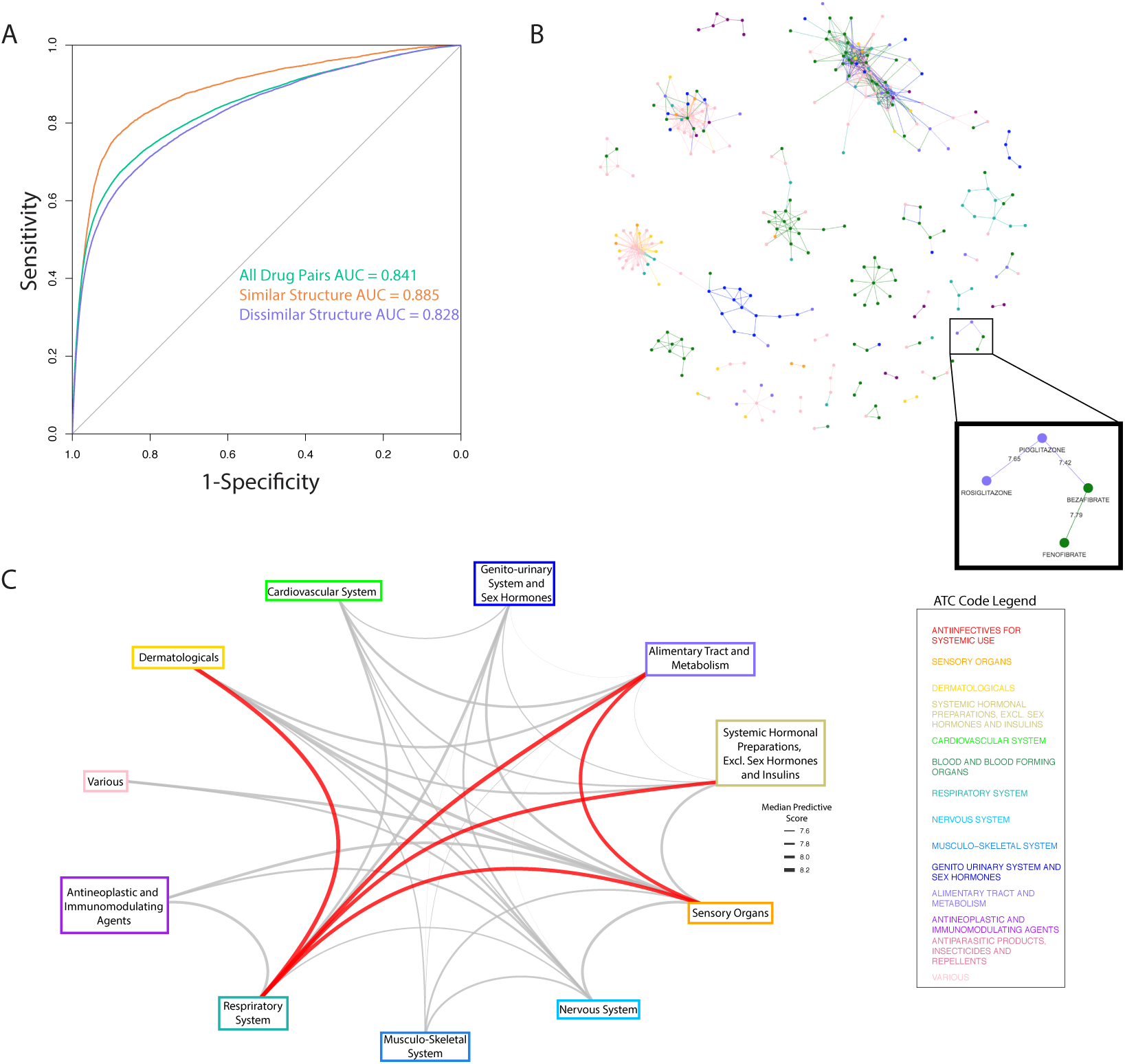
CATNIP model accurately predicts drugs that share an indication and can be used for repurposing. A) Receiver-operating characteristic curve for CATNIP, the performance for drug pairs with high and low structural similarity is also shown. B) A network of all drug pairs with a CATNIP score higher than 7.4. Nodes (drugs) are colored based on ATC classification and a specific example of repurposing between ATC classifications is highlighted. C) A graph of all ATC classification and the median CATNIP score between the drugs belonging to each of them (only including drug pairs with > 7.4 CATNIP score). The edges between ATC Classifications with the highest median CATNIP scores are colored red.

It has been shown before that structurally similar drugs have a high probability of treating the same indication^15^. However, there are many examples of drug pairs that defy this rule. For example, tamoxifen^16^ and anastrozole^17^ are structurally dissimilar compounds (Dice similarity = 0.372) that treat the same indication (Metathesaurus term: Cancer, Breast). To ensure that our model could accurately classify drug pairs that share an indication but are not structurally similar, we recalculated all performance metrics to control for high and low structural similarity. High performance was retained under both of these conditions (high structural similarity AUC = 0.885, low structural similarity AUC = 0.828 AUC, **Figure 2A**). These performance metrics confirm that our model is robust enough to predict if a drug pair will share an indication with or without structural similarity.

### Network clusters identify drugs with similar clinical characteristics

We constructed a repurposing network by calculating a CATNIP score for all possible drug pairs found within DrugBank, and assigning the drugs as nodes and the CATNIP score as the edge weight. We pruned the network using a cut-off value of 7.4 for the CATNIP scores (**Figure 2B**), which included 792 different drug pairs. This cut-off is equivalent to a predicted probability of >99% to share an indication and allowed for a balance between confidence within our predictions and drug diversity and availability.

We hypothesized that drugs sharing at least one indication would cluster together in our network. To confirm this theory, we classified each drug per its 1st order Anatomical Therapeutic Chemical (ATC) classification. This identification is a method of distinguishing the clinical use of a drug that is widely used in European and North American chemoinformatics databases^18^. Using ATC, we observed clearly defined clusters within the repurposing network (**Figure 2B**). Many clusters featured multiple ATC classifications, suggesting potential repurposing opportunities. For example, one cluster included the thiazolidinediones, rosiglitazone and pioglitazone (ATC classification: ‘Alimentary Tract and Metabolism’) and the fibrates, fenofibrate and bezafibrate (ATC classification: ‘Cardiovascular system’). These two clustered ATC classifications were connected by a high (7.42) CATNIP score between bezafibrate and pioglitazone, an antidiabetic drug; a relationship driven by the shared targeting of PPARa and PPARg resulting in the improvement of lipid and glucose metabolism. Bezafibrate has shown efficacy in the treatment of Type 2 Diabetes in numerous retrospective and pre-clinical studies, including Phase 2 trials^19–21^, however is still not an approved antidiabetic. The identification of bezafibrate as a potential diabetes treatment is a key example of how CATNIP can be used to identify repurposing opportunities.

We reasoned that the connections between ATC classifications across all the drug clusters could provide additional aid for drug repurposing purposes. Using the pruned network (CATNIP Score > 7.4), we collected all the scores between drugs of differing ATC classifications. From this collection, we were able to determine the median score associated between each pair of ATC classifications. The ATC classifications with the highest median CATNIP scores had literature support for numerous repurposing efforts between them (**Table 1**). For example, drugs with the ATC classifications of “Respiratory System” and “Systemic Hormonal Preparations, excluding sex hormones and insulins” were strongly connected to each other (7.97 median CATNIP score). This connection was driven by highly scored pairs of drugs including rimexolone to mometasone (8.31 CATNIP score) and prednisone to triamcinolone (8.13 CATNIP score). These connections are supported by the fact that hormonal agents like glucocorticoids and beta adrenergic agonists have been used for decades to relax the airway musculature in patients with reactive airways disease and chronic obstructive pulmonary disease^22^. Interestingly, our analysis identified glucagon, a peptide hormone that increases blood glucose levels, as a candidate for “Respiratory System” repurposing and this use already has clinical support^23,24^. Additionally, drugs classified as “Respiratory System” and “Dermatological” were also observed to be highly associated because of interactions such as the one between ciclesonide and hydrocortisone (8.43 CATNIP score). Ciclesonide and hydrocortisone do in fact share a clinical indication, “Asthma Bronchial”, giving added confidence to our findings. These types of network observations are important in laying the groundwork for suggesting novel clinical repurposing strategies for FDA-approved drugs.

**Table 1:** Literature Support for ATC Repurposing Predictions

| ATC Code 1 | ATC Code 2 | Reference |
| --- | --- | --- |
| Dermatologicals | Respiratory System | 64-68 |
| Alimentary Tract and Metabolism | Respiratory System | 69-72 |
| Sensory Organs | Respiratory System | 73-75 |
| Systemic Hormonal Preparations, Excluding Sex Hormones And Insulins | Respiratory System | 76, 77 |
| Sensory Organs | Alimentary Tract and Metabolism | 78-82 |

### CATNIP identifies novel disease areas for drug classes

We investigated the ability to leverage CATNIP scores to identify repurposing opportunities by evaluating specific drug classes. Drug classes are predefined in DrugBank. In order to identify actionable repurposing possibilities, we narrowed this list down to 50 classes containing inhibitors, antagonists, or agonists of specific gene or protein families. We focused our attention on specific disease areas that are attractive for drug repurposing opportunities, due to a lack of current treatments or high rates of acquired resistance. The specific disease areas were: “mental disorders”, “neurological diseases”, “diabetes”, and “cancer” (cancer was further divided into specific cancer types due to the large variance in disease pathology between types, **Methods**). We hypothesized that CATNIP scores could be used to identify specific drug classes that would be efficacious for a new disease area. For each drug class and disease area, we found the statistical difference in the CATNIP score distribution between two sets of drug pairs. The first set included pairs that had one drug within the drug class and the other drug approved for the disease in question, while the other set included drug pairs that had one drug within the drug class and the other drug not approved for the disease in question (**Methods**). We compared the effect size, estimated by the Wilcoxon location shift, for all drug class-disease pairs that had a significant difference in distribution compared to drug class-non-disease pairs (FDR < 0.1, **Figure 3A-B, Figure S7-8**). By using CATNIP scores, we found that many well-known drug class-diseases associations could be recovered. For example, “muscarinic antagonists” were highly ranked for “neurological diseases” and many such agents are FDA-approved for this indication^25^. In addition, we found that “kinase inhibitors” were closely associated with the treatment of cancer and “dopamine antagonists” for the treatment of “mental disorders”^26, 27^ (Wilcoxon Location Shift = 0.711-0.945 for “kinase inhibitors” and select cancer types, Location Shift = 0. 882 for “dopamine antagonists” and “mental disorders”, p-value < 0.001, **Figure S9**). In fact, almost all drug class-disease associations contained at least one FDA-approved drug for the respective disease, giving us added confidence in our model. Of note, each drug was allowed to be categorized into numerous drug classes, leading to unexpected, yet easily explained, results; for example, “dopamine antagonists” appearing as a top drug class for “neurological diseases”. This is due to risperidone, a drug traditionally used for schizophrenia and mood disorders, also having a secondary indication of Alzheimer’s type severe dementia.

**Figure 3:**
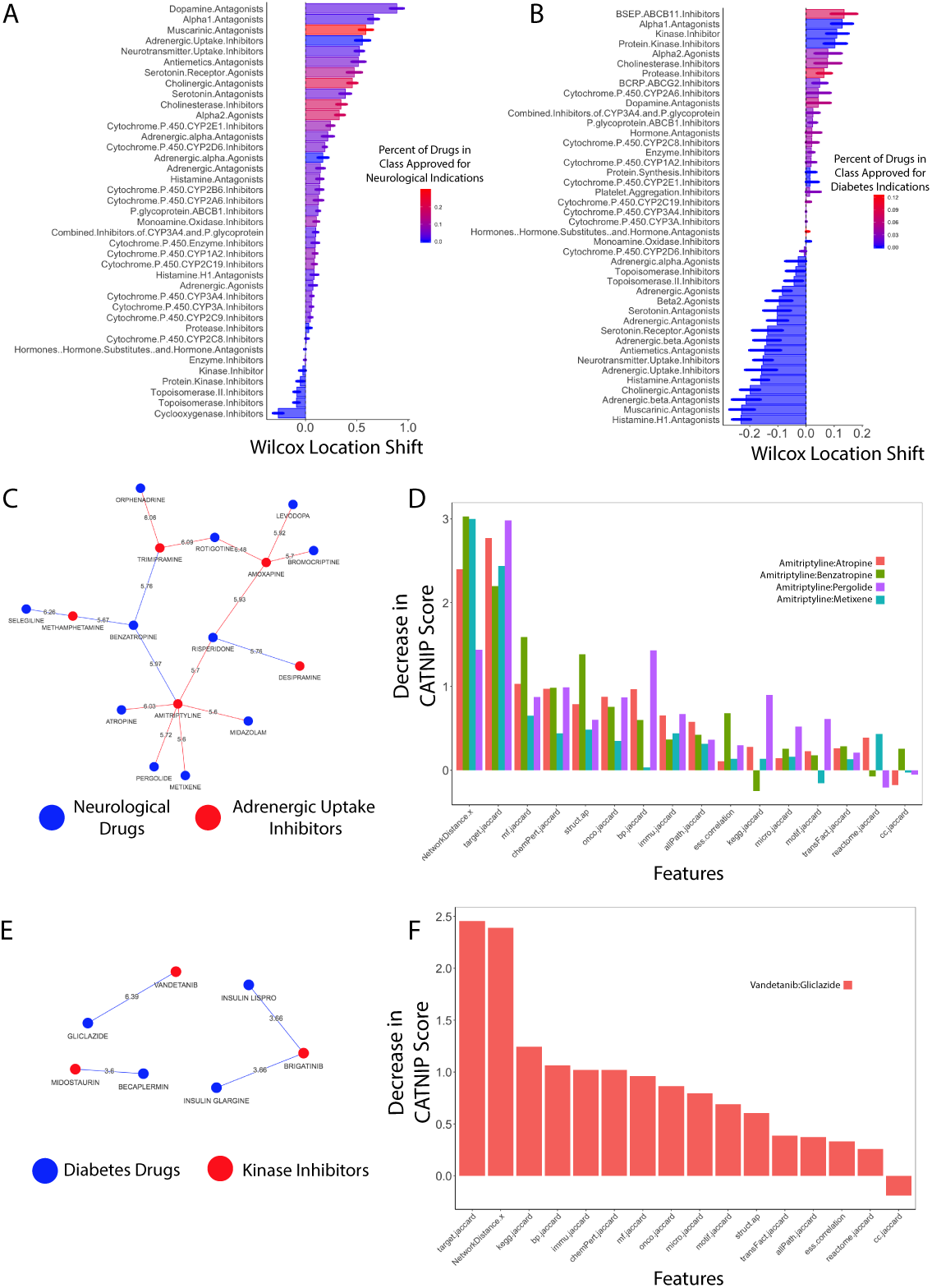
CATNIP identifies drug class repurposing opportunities. The location shift, calculated using Wilcox-Mann-Whitney for all CATNIP scores of drug class-disease drug pairs vs drug class-non-disease drug pairs for A) “neurological diseases” and B) “diabetes”. Drug classes are colored based on the percent of drugs within the class that are approved for treatment of the specific disease and only significant associations are shown (FDR < 0.1). C) The network of neurological drugs and adrenergic uptake inhibitors drug pairs with the highest CATNIP scores. D) The decrease in the CATNIP score when removing each feature for amitriptyline and select Parkinson’s Disease drugs. E) The network of anti-diabetes and kinase inhibitor drug pairs with the highest CATNIP scores. F) The decrease in the CATNIP score when removing each feature for the drug pair vandetanib and gliclazide.

Next, we further interrogated the drug classes associated with “neurological diseases” and “diabetes”, specifically. CATNIP scores were able to correctly identify almost all drug classes known to treat these diseases (**Figure 3A-B**). To identify possible repurposing candidates, we focused our attention on drug classes shown to have a large positive effect size with this CATNIP analysis but are not currently approved for treatment. For “neurological diseases”, the use of adrenergic uptake inhibitors, traditionally used as antidepressants, was the top repurposing candidate (**Figure 3A**). For “diabetes” alpha 1 antagonists and kinase inhibitors were identified as possible novel treatments for diabetes (**Figure 3B**). We believe further investigation into these drug classes and diseases could lead to successful clinical applications.

### CATNIP interpretability reveals reasoning for repurposing candidates

From our list of repurposing candidates, we chose two novel drug class-disease associations to further investigate.

#### Adrenergic uptake inhibitors applied to Parkinson’s disease

First, we evaluated the relationship between “neurological diseases” and “adrenergic uptake inhibitors”. We focused on the drug pairs with the highest CATNIP scores, i.e. those predicted with the highest confidence to share at least one indication (**Figure 3C**). Of all the adrenergic uptake inhibitors, we found that amitriptyline and trimipramine, two anti-depressants, had the highest CATNIP scores with the “neurological diseases” drugs. The drugs that shared the strongest connections with amitriptyline and trimipramine were drugs approved for Parkinson’s disease (PD). Specifically, metixene, atropine, pergolide and benzatropine were associated with amitriptyline, according to CATNIP, and trimipramine was associated to benzatropine and rotigotine. Trimipramine was also strongly connected with orphenadrine, which is sometimes used off label in PD, but will not be included in the following analyses.

Using the CATNIP model, we evaluated which features contributed towards the prediction of amitriptyline and trimipramine to share an indication with PD drugs. We found that target, gene ontology, and pathway similarity all strongly contributed to the predictions for both amitriptyline and trimipramine (**Figure 3D, Figure S10**). Since target similarity and distance between targets (in a protein-protein interaction network) were among the top contributing features, we investigated which gene targets were shared amongst these drug pairs. We found that amitriptyline targets three specific gene classes that are also targeted by at least one of the PD drugs: muscarinic acetylcholine receptors, G-coupled protein receptors (GPCRs), and alpha adrenergic receptor. Trimipramine also targets muscarinic acetylcholine receptors, alpha-adrenergic receptors, and dopamine transporters, which is similar to benzatropine, a PD drug. All these receptors have well-defined relationships with PD and other neurological diseases^25, 28, 29^, which adds support for repurposing amitriptyline and/or trimipramine.

Amitriptyline may be an ideal candidate for use in PD patients. We evaluated the shared molecular function gene ontology terms shared between amitriptyline and all four PD drugs. GPCR activity was once again identified (**Supplementary Data**). We then interrogated the biological pathways these drug targets are involved in and found many broad GPCR pathways overlapping between amitriptyline and the PD drugs (**Figure S11**) including the Reactome pathway “GASTRIN_CREB_SIGNALLING PATHWAY VIA PKC AND MAPK”. Several recent studies support the link between gastrin-releasing peptide signaling to brain function^30^. Through CATNIP, we have identified “adrenergic uptake inhibitors” like amitriptyline and trimipramine as a possible treatment for PD.

#### Kinase inhibitors applied to Diabetes

Our CATNIP analysis identified an opportunity to repurpose “kinase inhibitors” for the treatment of diabetes (**Figure 3B**). Of the drug pairs evaluated in this context, the link between vandetanib, a thyroid cancer drug, and gliclazide, a Type 2 diabetes drug (CATNIP Score = 6.39, **Figure 3E**) was the strongest. This association was driven by target similarity and similarity between KEGG pathways of the drug targets (**Figure 3F**). Vendetanib and gliclazide have an overlapping target, VEGFA. Several KEGG pathways are shared between vandetanib and gliclazide including the “Cytokine cytokine receptor interaction” pathway (**Supplementary Data)**. This pathway contains VEGFA, the shared target, and the epidermal growth factor receptor (EGFR), another one of vandetanib’s targets. The similarity between these two drug’s targets and pathway effects leads us to believe there is strong potential for vendetanib to be repurposed.

## Discussion

Although considerable improvements have been made in drug repurposing efforts over the past decade, the use of previous disease associations will eventually curtail these improvements due to the imposed restriction of previous knowledge. Our new approach, CATNIP, could provide a highly effective aid to drug repurposing endeavors. Here, we accurately predicted drugs that shared an indication, while keeping high levels of both sensitivity and specificity. Leveraging our prediction metric enabled us to generate a network for repurposing, identifying, and repurposing predictions based on system-wide drug scopes.

The CATNIP method allows for broad-scale drug repurposing opportunities to be readily identified. By identifying and interpreting our drug similarity features, we can investigate the possible mechanisms behind these repurposing candidates. The benefit of using drug similarity features is two-fold. First, these features are readily available for both approved and investigational drugs, which have been underserved by previous repurposing methods. Second, the interpretability of the features allows us to identify possible mechanisms of action when we back engineer what contributed to high CATNIP scores.

We found strong support for repurposing amitriptyline and trimipramine, both of which are in clinical use as anti-depressants, for PD. These drugs have many functions in addition to being adrenergic uptake inhibitors, such as serotonin blockers, anticholinergics, and the mechanisms overlapping with current PD drugs described above. Movement Disorders Society guidelines found insufficient evidence to support the use of amitriptyline for depression in PD^31^ and a published Practice Parameter found only level C evidence for its use^32^. However, amitriptyline has been commonly used for not only depression but other off-label indications in neurological disorders, including pain^33^. While clinical trials have been conducted for the effect of amitriptyline on depression in PD patients^34^, currently there are no trials evaluating amitriptyline or trimipramine as a treatment for other symptoms and signs of PD. There have, however, been preclinical studies evaluating amitriptyline as a potential therapy for PD. In rodent models of PD, amitriptyline affects levels of neurotrophic factors including BDNF^35^ and decreases dopamine cell loss in these models^36, 37^. It has been suggested to mitigate microglial inflammation^38^. Moreover, with the suggestion that amitriptyline may have shorter term symptomatic motor benefit, it may enhance levodopa efficacy^39^.

When we more closely evaluated trimipramine, we found compelling evidence this could be a potential PD therapeutic. Specifically, the targets of trimipramine make it a potentially strong therapeutic to combat loss of motor function amongst PD patients. This benefit is due to the dual targeting of DRD2 and alpha 2 adrenergic receptors, which is similar to piribedil, an investigational PD medication that was not included within our final CATNIP network due to a lack of available information. In a review of piribedil, it was highlighted that the agonistic D2/D3 activity combined with alpha 2 adrenergic antagonism can lead to preservation of motor function^40^. However, further research must be done to better understand the exact effects that trimipramine has on both dopamine and alpha 2 adrenergic receptors. Further research into trimipramine could quickly lead to a clinical trial for PD patients with specific motor function end points.

We also identified a repurposing opportunity with kinase inhibitors for the treatment of diabetes, due to the strong predicted connection between vandetanib, a thyroid cancer drug, and gliclazide. While there have been some preclinical animal studies investigating the use of kinase inhibitors in diabetes^41, 42^, to our knowledge, there has yet to be an approved kinase inhibitor for the treatment of diabetes. Both vandetanib and gliclazide are known to target VEGFA, which has shown a clear connection to diabetes pathology^43^ and treatment^44^. Additionally, Hagberg et al. published work suggesting that antagonism of VEGFB, a gene within the same pathway as VEGFA, improves insulin sensitivity and increases skeletal muscle glucose uptake in *db/db* mice^45^. Because vandetanib targets VEGFR1^46^, the receptor VEGFB binds, it could also have insulin sensitizing effects. Further experimental work is required to verify this hypothesis^47^.

Besides the targeting of VEGFA/VEGFR1, vandetanib’s target EGFR can also potentially help diabetes pathology. Inflammatory cytokines (including, but not limited to, IL-8 and TNF-α) have been shown to be associated with the progression of diabetic neuropathy^48^. The inhibition of EGFR through the use of a kinase inhibitor in past work has reduced the expression of both to IL-8 and TNF-α in rats^49^. Therefore, we believe vandetanib could be considered as a potential diabetes treatment, due to its ability to target EGFR leading to a possible decrease in inflammatory cytokine production.

In addition to the exciting predicted repurposing opportunities we have chosen to highlight, many other drug classes showed significant repurposing potential for mental disorders, neurological diseases, and several different cancer types. While diving into each of these opportunities is outside the scope of this paper, we hope that researchers take it upon themselves to further investigate these candidate drug class-disease associations.

It is important to acknowledge certain limitations to CATNIP, such as data availability and the application to rare diseases. Although this model does not rely on disease similarity information, it does require known molecular target information to obtain peak predictive power. This target information can frequently be unavailable for early stage compounds. Additionally, this method would have limited use when searching for drugs to be repurposed for diseases with very few or no clinically approved compounds.

To our knowledge, CATNIP is the first method capable of predicting a novel indication for a drug without relying on disease similarities. Many predictions gained from CATNIP have substantial preclinical research or mechanistic support, suggesting that other predictions may also provide valuable information for future investigations. Due to its demonstrated ability to identify large scale drug repurposing opportunities, CATNIP will likely serve as a significant basis towards a bright future in drug repurposing efforts.

## Methods

### Indication Mapping

Using a custom Python script, we webscraped DrugBank 5.0^50^ for drug compound names and indication information with a total of 3066 drugs being found. Indication information were run through the Unified Medical Language System (UMLS) tool, MetaMap^13^, to match DrugBank assigned indications to MESH IDs and UMLS Concept Unique Identifiers (CUIs). MetaMap is a computational approach that combines linguistic and natural language processing techniques to map biomedical texts to the UMLS Metathesaurus. MetaMap has previously been shown to successfully exceeded human mapping capabilities^14^. Using a custom Python script we identified synonym candidate to further improve indication semantics. A random subset of the indications were manually reviewed and found to correctly map to standardized terms with a 95% accuracy (**Figure S1B**). We then filtered our list of drugs to the 2576 drugs that shared at least one indication with another drug.

### Similarity Feature Collection

#### Compound Features

Similarities between drugs were found by creating all possible pairs of the drugs with known indications. Multiple compound similarity features and drug target similarity features were collected. The drug targets listed within DrugBank 5.0^50^ were used as our set of ‘known targets’ for each drug. Additionally, we collected genomic information about each drug target using MSigDB^51, 52^. The features, sources and metrics used to measure similarity are listed in **Supplementary Table 1**. The measures of similarity included, but were not limited to, Pearson Correlation, Jaccard Index, and Dice Similarity. In cases where there was insufficient or missing information, features were imputed by using the median value for that feature in drug pairs with complete information.

#### Network Features

We curated a biological network that contains 22,399 protein-coding genes, 6,679 drugs, and 170 TFs. The protein-protein interactions represent established interaction^53–55^, which include both physical (protein-protein) and non-physical (phosphorylation, metabolic, signaling, and regulatory) interactions. The drug-protein interactions were curated from several drug target databases^55^.

### Statistical Analysis

For each similarity feature, a Kolmogorov-Smirnov (KS) test was performed between the set of drug pairs that shared an indication and those that did not share an indication. The KS test was chosen to identify non-linear predictive power. In addition, the Pearson correlation between all numeric features was calculated. These tests were performed using custom scripts in R statistical software ^56^.

### Model Building

We trained a two-class classifier predictive model using the features described above. Our classes were determined as a binary of “shared” or “non-shared” indication. Drugs were only included if they shared an indication with at least one other drug. A 5-fold cross-validation gradient boosting model was used after careful model selection and implemented using the XGBoost package^57^ within the R statistical software. Additional models that were tested and compared using the AUC and AUPRC of 5-fold cross-validation were: Support Vector Machine with a radial kernel model, logistic regression with elastic net and logistic regression with lasso, all using custom R scripts. A custom-made R script was used to implement a grid-search to optimize the hyperparameters of our model. Our model objective was a logistic regression for binary classification and we output a score pre-logistic transformation. The class size of “shared” vs. “non-shared” was imbalanced, therefore we applied downsampling to each fold of training via the R package Caret^58^.

### Classification Evaluation

For evaluating the model performance on predicting if two drugs share an indication, receiver operating characteristic (ROC) and precision-recall curve (PRC) curves were created in R using the pROC^59^ and precrec^60^ packages respectively. The raw-logistic values were normalized on a scale from 0-1 to enable easier interpretation and ROC/PRC calculation. Area-under-the-ROC curve (AUC) and area-under-the-PRC (AUPRC) scores were used to evaluate model performance.

### Drug Similarity Network

#### Network Construction

We constructed a drug similarity network based upon our classifier results with drugs as nodes and our raw model output as the edge weight. This network was visualized using the visNetwork package^61^ and used in analyses using the iGraph package^62^ within R^56^.

#### ATC Repurposing Analysis

The Anatomical Therapeutic Chemical (ATC) code for all drugs were found in DrugBank^50^, and the highest level code was assigned. Drugs with multiple ATC codes assigned to them were re-assigned into the category “Various”. A circular repurposing network was created with ATC codes as the nodes using the iGraph^62^ and gGraph^63^ packages with R^56^. The graph edge weights were based on the mean classifier output between all drugs of each ATC code category. To reduce noise within the repurposing network an initial cut-off of drug pairs with a classifier output of 7.4 and above was implemented, leaving 792 drug pairs to examine. Manual literature searches were used to validate repurposing opportunities.

#### Drug Class Repurposing Analysis

Drug classes for all drugs were found in DrugBank^50^ and were filtered to include only “inhibitor”, “antagonist,” and “agonist” classes that had at least 20 drugs, to ensure enough statistical power. Additionally, we identified four main disease areas of interest: “mental disorders”, “neurological diseases”, “diabetes”, and “cancer”. The UMLS^13^ sematic codes “modb” and “neop” were used to identify indications falling within mental disorders and cancer, respectively. Cancer was further refined into different cancer types based on a keyword search in a custom Python script. All UMLS concept IDs containing the word “diabetes” were included within the diabetes category. For “neurological diseases”, we refined our list to only include Parkinson’s Disease, Alzheimer’s, Epilepsy, and Dementia, to balance both specificity in disease type and enough drugs to make statistically sufficient sample size.

Wilcox-Mann-Whitney tests between all drug class-disease associations were performed. The test specifically tested if the mean of the CATNIP scores of drug pairs with one drug being a member of the class of interest and the other being approved for the disease of interest were significantly different than the mean of the CATNIP scores of all drug pairs that included one drug within the class of interest and the other drug not being approved for the diseases of interest. A positive location shift meant that drug class-disease pairs had significantly higher CATNIP scores than drug class-non-disease pairs. A FDR multiple hypothesis correction was applied.

#### CATNIP Feature Effect Analysis

The effect of each feature on the CATNIP score for specific drug pairs was found by iteratively changing the feature value to the median value of that feature for all drug pairs. Since the clear majority of all drug pairs do not share an indication this is the best approximate for that feature having no contribution to the CATNIP score. The difference in the new CATNIP score and the correct CATNIP score was then measured.

## Acknowledgements

C.M.G., J.E, N.S.M., and O.E. conceived, designed and developed the methodology for this work. C.M.G., J.E., and O.E. analyzed and interpreted the data. C.M.G. executed the machine learning analyses. C.M.G. and J.E. wrote the initial draft of the manuscript. M.D.G. provided expertise within diabetes, as well as other clinical aspects of the paper. C.H. provided expertise within Parkinson’s Disease, as well as other clinical aspects of the paper. O.E. supervised the study. All the authors reviewed and approved the manuscript.

## Supplementary Figures

Figure S1: MetaMap performs well in drug indication mapping. A) The number of occurences of different UMLS sematic types. B) The accuracy of mapping indications using MetaMap for indications categorized a “Structured” and the “Description” section.

Figure S2: Target ontology similarity data types vary for drug pairs that share an indication and those that do not. The violin plots of similarity distributions for the similarities of targets’ A) biological processes, B) cellular component, C) molecular function, D) chemical perturbation, E) oncological, F) immunogenic, G) micro-RNA, and H) transcription factor. Statistical significance found by Kolmogorov-Smirnov test.

Figure S3: Target similarity data types vary for drug pairs that share an indication and those that do not. The violin plots of similarity distributions for the similarities of A) targets, B) the Protein-Protein Interaction network distance between targets and the C) correlation of target essential within cancer cell lines. Statistical significance found by Kolmogorov-Smirnov test.

Figure S4: Target pathway similarity data types vary for drug pairs that share an indication and those that do not. The violin plots of similarity distributions for the similarities of the A) reactome pathways, B) all pathway types and C) KEGG pathways a drug’s target is known to be involved within. Statistical significance found by Kolmogorov-Smirnov test.

Figure S5: Structure similarity varies for drug pairs that share an indication and those that do not. A) The violin plot of the Dice chemical fingerprint similarity, statistical significance found by Kolmogorov-Smirnov test.

Figure S6: CATNIP performs significantly better than random. A) The Precision – Recall curve for classifying if two drugs share an indication using CATNIP and the random expectation.

Figure S7: CATNIP identifies drug class repurposing opportunities. The location shift, calculated using Wilcox-Mann-Whitney for all CATNIP scores of drug class-disease drug pairs vs drug class-non-disease drug pairs for A) Mental Disorder, B) Skin Cancer, C) Lung Cancer, D) Breast Cancer, E) Thyroid Cancer, F) Large Intestine Cancer, G) Upper Aerodigestive Tract Cancer H) Gastric Cancer I) Renal Cancer. Drug classes are colored based on the percent of drugs within the class that are approved for treatment of the specific disease and only significant associations are shown (FDR < 0.1).

Figure S8: CATNIP identifies other drug class repurposing opportunities within cancer. The location shift, calculated using Wilcox-Mann-Whitney for all CATNIP scores of drug class-disease drug pairs vs drug class-non-disease drug pairs for A) Urinary Tract Cancer, B) Pleura Cancer, C) Endometrium Cancer, D) Ovarian Cancer, E) Pancreatic Cancer, F) Bone Cancer, G) Oesophagus Cancer, H) Lymphoma/Leukemia, and I) Autonomic Ganglia Cancer. Drug classes are colored based on the percent of drugs within the class that are approved for treatment of the specific disease and only significant associations are shown (FDR < 0.1).

Figure S9: CATNIP scores are statistically higher between drugs of certain drug classes and drugs that treat associated diseases. The distributions of CATNIP score between A) kinase inhibitors and drugs known to treat cancer and those that do not and B) dopamine antagonists and drugs known to treat mental illness and those that do not.

Figure S10: Target features drive the prediction of trimipramine as a Parkinson’s Disease treatment. A) The decrease in the CATNIP score when removing each feature for trimipramine and select Parkinson’s Disease drugs.

Figure S11: Many pathways or gene ontology groups overlap, fueling CATNIP predictions. The overlap between amitriptyline and select Parkinson’s Disease drugs for A) reactome pathways, B) KEGG pathways, and D) molecular function gene ontologies. The overlap between vandetanib and gliclazide for A) reactome pathways, B) KEGG pathways, and D) molecular function gene ontologies.

## Supplementary Tables

Table S1: The drug similarity features used within CATNIP.

Table S2: Comparison of model performance using other model types.

## Supplementary Data

File 1: All pathways and gene ontologies that amitriptyline’s targets and the targets of select Parkinson’s Disease drugs’ targets are associated with.

File 2: All pathways and gene ontologies that trimipramine’s targets and the targets of select Parkinson’s Disease drugs’ targets are associated with.

File 3: All pathways and gene ontologies that vandetanib’s targets and gliclazide’s are associated with.

