## Supplemental Figures for "A machine learning and network framework to discover new indications for small molecules"

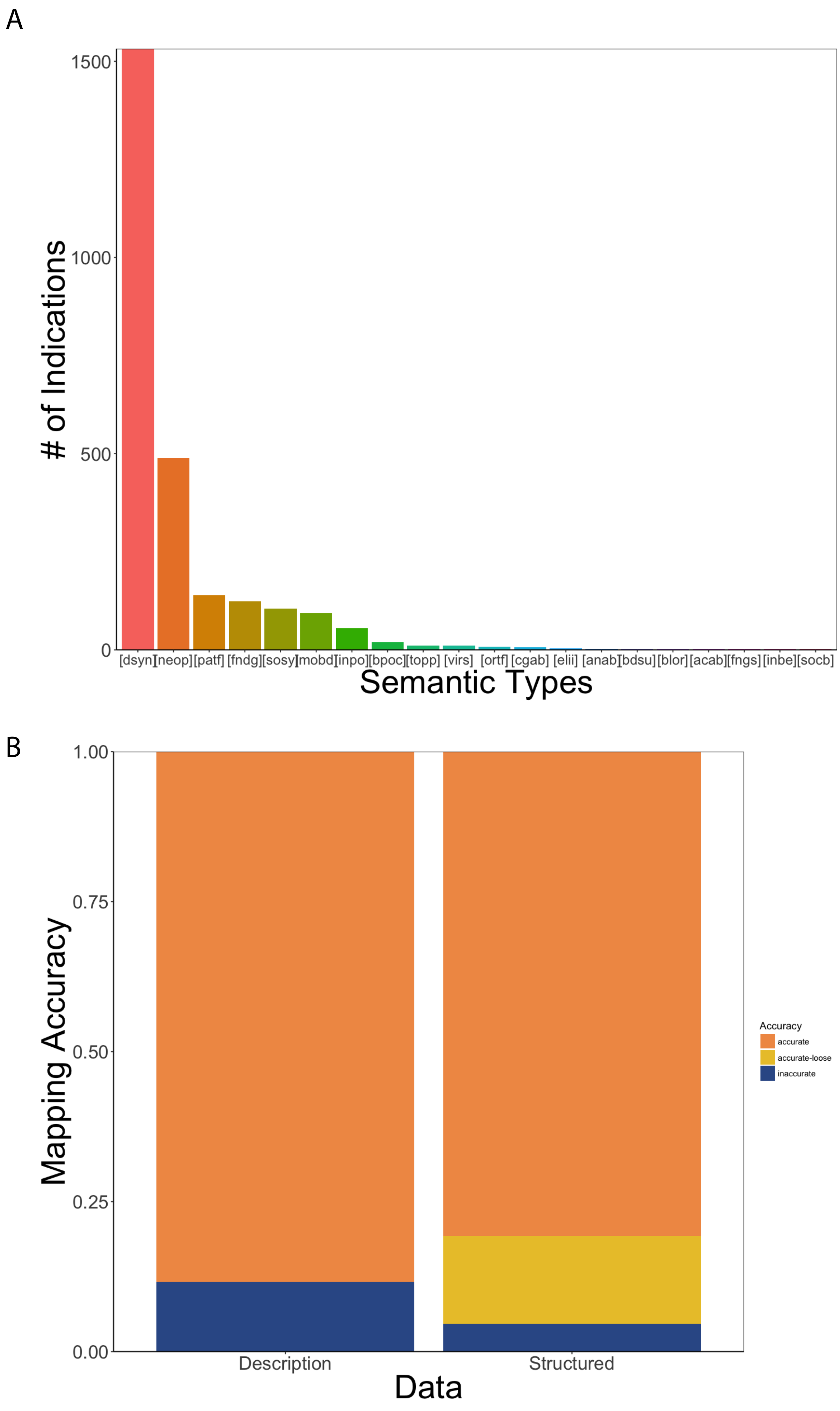
A
1500
1000
500
0
### of Indications
[dsyn] [neop] [patf] [fndg] [sosy] [mobd] [inpo] [bpoc] [topp] [virs] [ortf] [cgab] [elii] [anab] [bdsu] [blor] [acab] [fngs] [inbe] [socb]
Semantic Types
B
1.00
0.75
0.50
0.25
0.00
Mapping Accuracy
Description
Structured
Data
Accuracy
accurate
accurate-loose
inaccurate

Figure S1

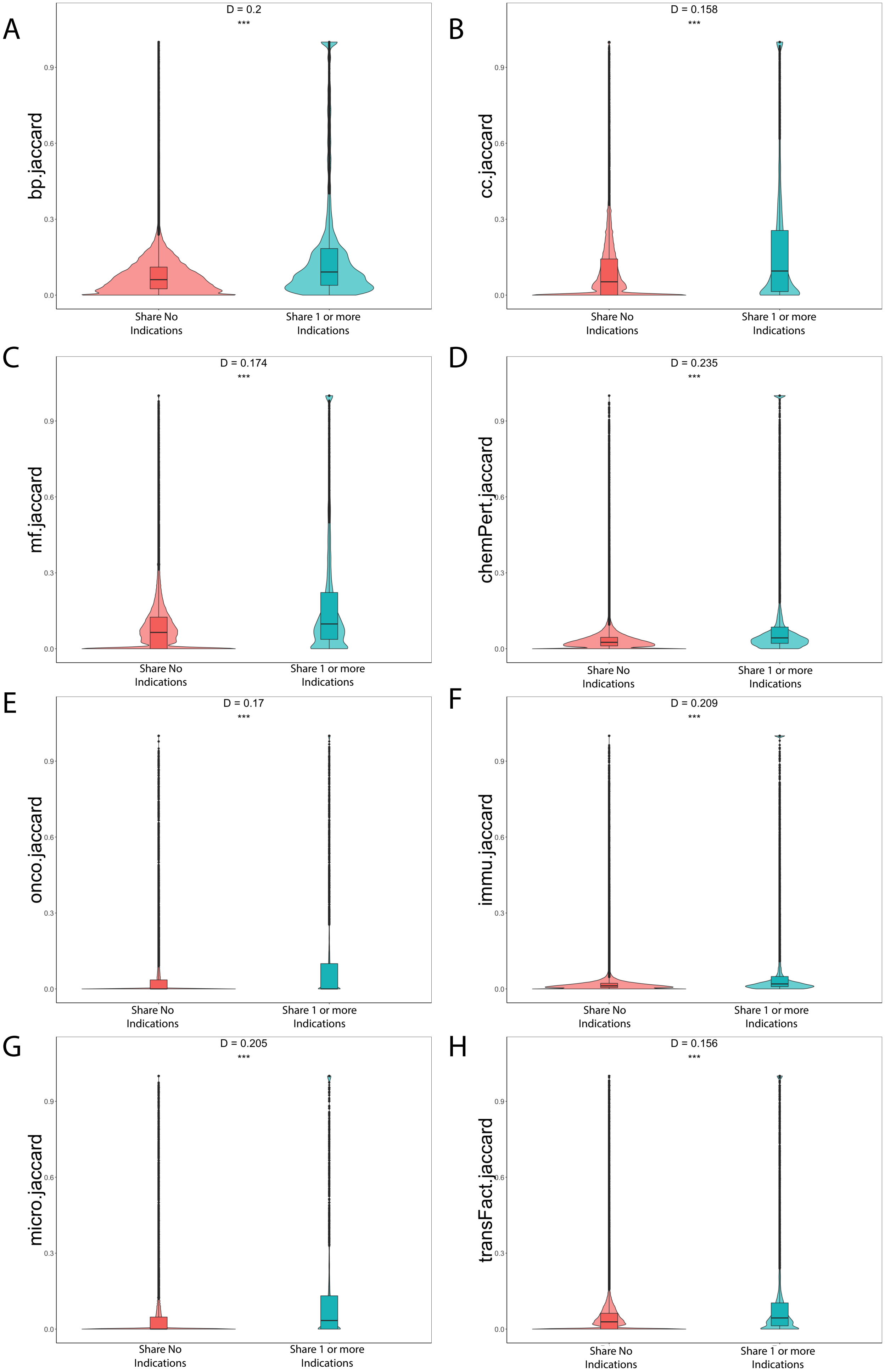
A
D = 0.2
***
bp.jaccard
0.0
0.3
0.6
0.9
Share No Indications
Share 1 or more Indications
B
D = 0.158
***
cc.jaccard
C
D = 0.174
***
mf.jaccard
D
D = 0.235
***
chemPert.jaccard
E
D = 0.17
***
onco.jaccard
F
D = 0.209
***
immu.jaccard
G
D = 0.205
***
micro.jaccard
H
D = 0.156
***
transFact.jaccard

Figure S2

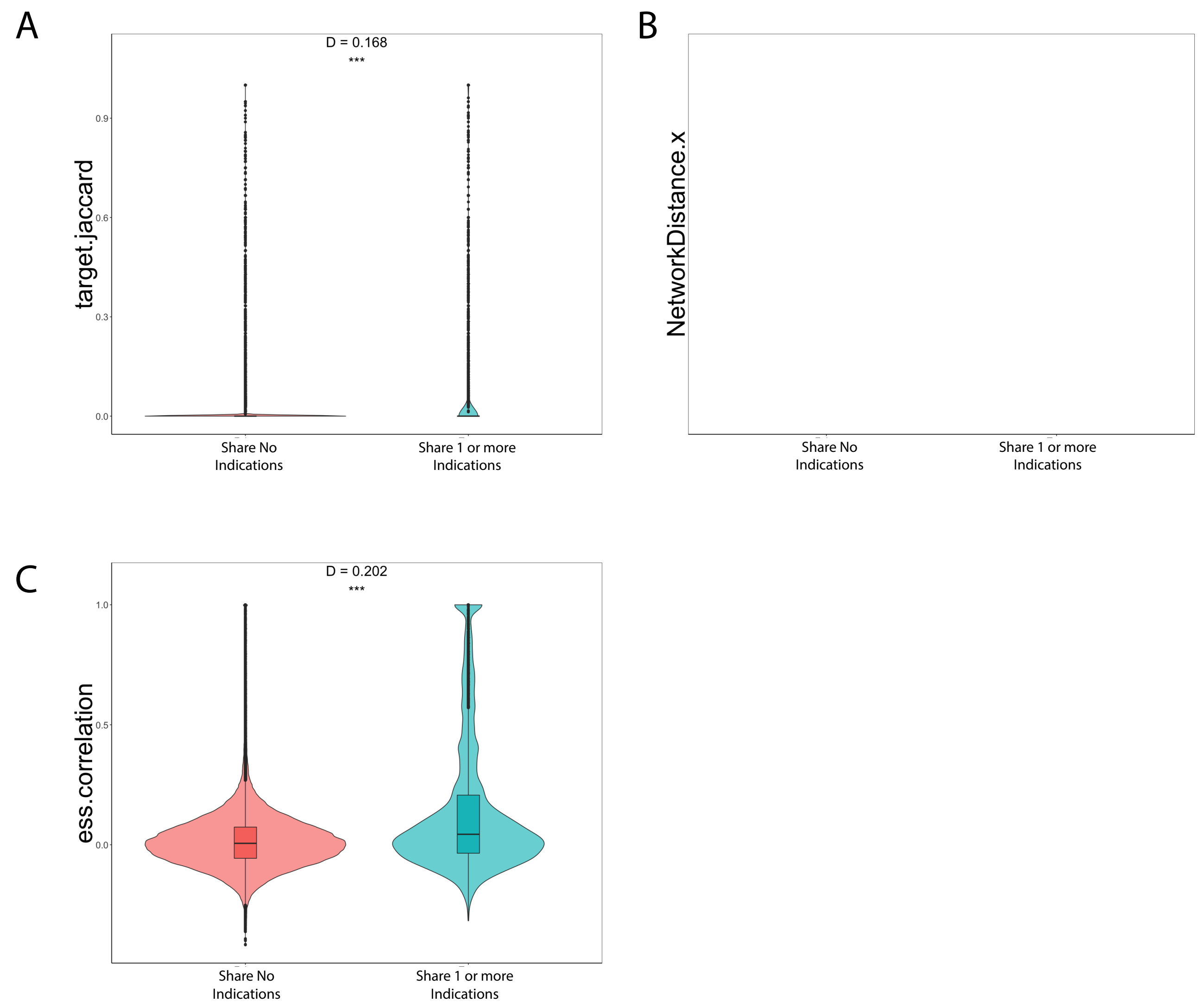
A
D = 0.168
***
target.jaccard
0.9
0.6
0.3
0.0
Share No Indications
Share 1 or more Indications
B
NetworkDistance.x
Share No Indications
Share 1 or more Indications
C
D = 0.202
***
ess.correlation
1.0
0.5
0.0
Share No Indications
Share 1 or more Indications

Figure S3

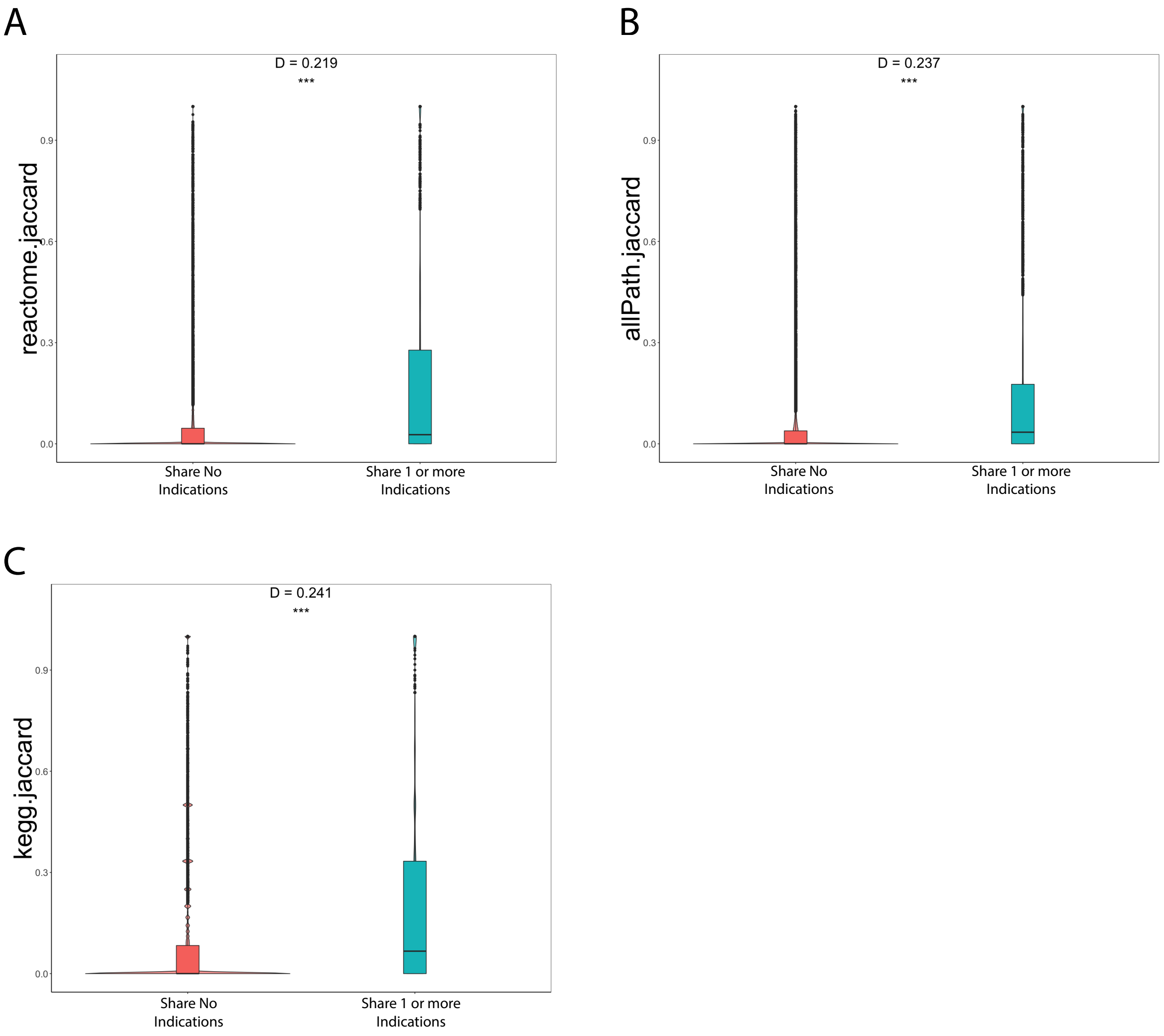
A
D = 0.219
***
reactome.jaccard
0.0
0.3
0.6
0.9
Share No Indications
Share 1 or more Indications
B
D = 0.237
***
allPath.jaccard
0.0
0.3
0.6
0.9
Share No Indications
Share 1 or more Indications
C
D = 0.241
***
kegg.jaccard
0.0
0.3
0.6
0.9
Share No Indications
Share 1 or more Indications

Figure S4

A

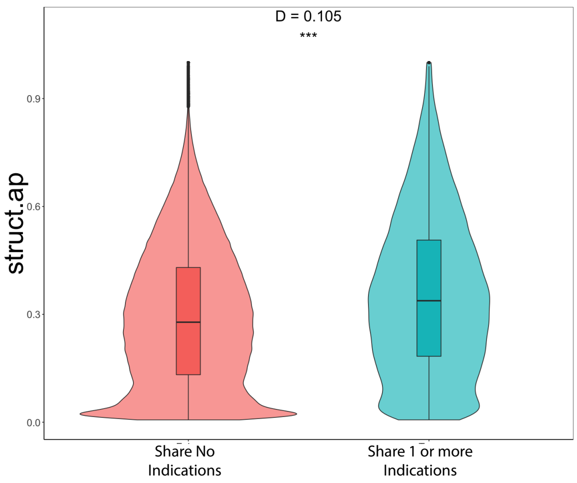
D = 0.105
***
struct.ap
0.9
0.6
0.3
0.0
Share No
Indications
Share 1 or more
Indications

Figure S5

A

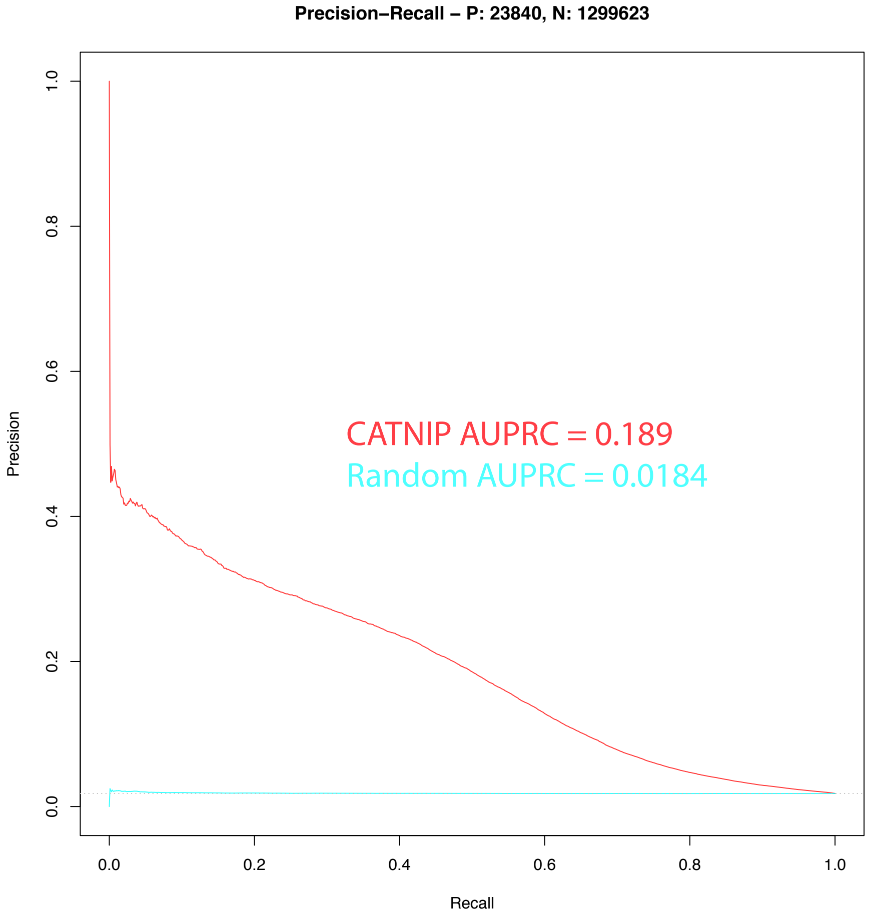

Precision−Recall − P: 23840, N: 1299623
Precision
1.0
0.8
0.6
0.4
0.2
0.0
CATNIP AUPRC = 0.189
Random AUPRC = 0.0184
0.0
0.2
0.4
0.6
0.8
1.0
Recall


Figure S6

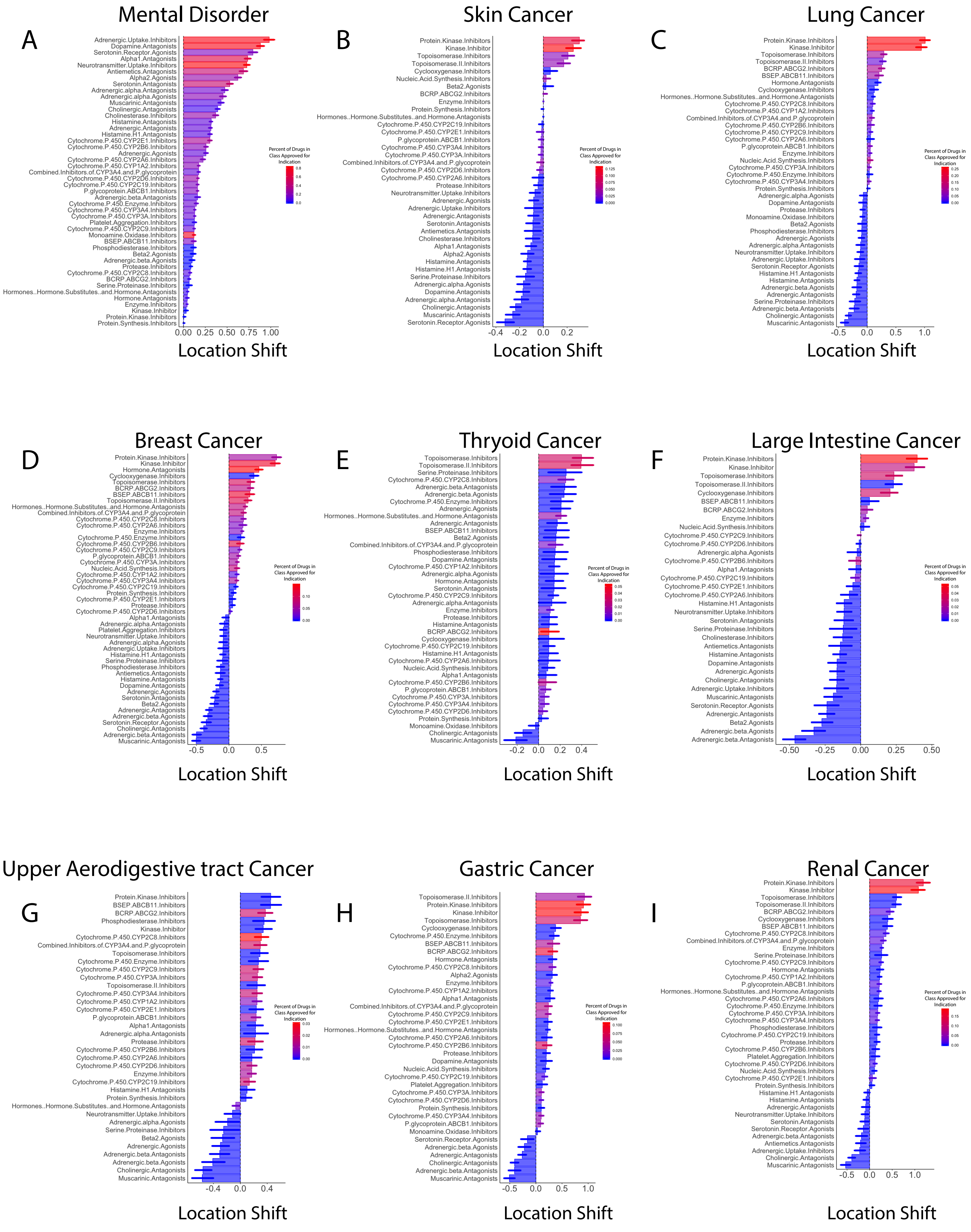
Mental Disorder
A
Adrenergic.Uptake.Inhibitors
Dopamine.Antagonists
Serotonin.Receptor.Agonists
Alpha1.Antagonists
Neurotransmitter.Uptake.Inhibitors
Antiemetics.Antagonists
Alpha2.Agonists
Serotonin.Antagonists
Adrenergic.alpha.Antagonists
Adrenergic.alpha.Agonists
Muscarinic.Antagonists
Cholinergic.Antagonists
Cholinesterase.Inhibitors
Histamine.Antagonists
Adrenergic.Antagonists
Histamine.H1.Antagonists
Cytochrome.P.450.CYP2E1.Inhibitors
Cytochrome.P.450.CYP2B6.Inhibitors
Adrenergic.Agonists
Cytochrome.P.450.CYP2A6.Inhibitors
Cytochrome.P.450.CYP1A2.Inhibitors
Combined.Inhibitors.of.CYP3A4.and.P.glycoprotein
Cytochrome.P.450.CYP2D6.Inhibitors
Cytochrome.P.450.CYP2C9.Inhibitors
P.glycoprotein.ABCB1.Inhibitors
Adrenergic.beta.Antagonists
Cytochrome.P.450.Enzyme.Inhibitors
Cytochrome.P.450.CYP3A4.Inhibitors
Cytochrome.P.450.CYP3A.Inhibitors
Platelet.Aggregation.Inhibitors
Cytochrome.P.450.CYP2C19.Inhibitors
Monoamine.Oxidase.Inhibitors
BSEP.ABCB11.Inhibitors
Phosphodiesterase.Inhibitors
Beta2.Agonists
Adrenergic.beta.Agonists
Protease.Inhibitors
Cytochrome.P.450.CYP2C8.Inhibitors
BCRP.ABCG2.Inhibitors
Serine.Proteinase.Inhibitors
Hormones..Hormone.Substitutes..and.Hormone.Antagonists
Hormone.Antagonists
Enzyme.Inhibitors
Kinase.Inhibitor
Protein.Kinase.Inhibitors
Protein.Synthesis.Inhibitors
0.00 0.25 0.50 0.75 1.00
Location Shift
Percent of Drugs in Class Approved for Indication
0.8
0.6
0.4
0.2
0.0
Skin Cancer
B
Protein.Kinase.Inhibitors
Kinase.Inhibitor
Topoisomerase.Inhibitors
Topoisomerase.II.Inhibitors
Cyclooxygenase.Inhibitors
Nucleic.Acid.Synthesis.Inhibitors
Beta2.Agonists
BCRP.ABCG2.Inhibitors
Enzyme.Inhibitors
Protein.Synthesis.Inhibitors
Hormones..Hormone.Substitutes..and.Hormone.Antagonists
Cytochrome.P.450.CYP2C19.Inhibitors
Cytochrome.P.450.CYP2E1.Inhibitors
P.glycoprotein.ABCB1.Inhibitors
Cytochrome.P.450.CYP3A4.Inhibitors
Cytochrome.P.450.CYP3A.Inhibitors
Combined.Inhibitors.of.CYP3A4.and.P.glycoprotein
Cytochrome.P.450.CYP2D6.Inhibitors
Cytochrome.P.450.CYP2A6.Inhibitors
Protease.Inhibitors
Neurotransmitter.Uptake.Inhibitors
Adrenergic.Agonists
Adrenergic.Uptake.Inhibitors
Adrenergic.Antagonists
Serotonin.Antagonists
Antiemetics.Antagonists
Cholinesterase.Inhibitors
Alpha1.Antagonists
Alpha2.Agonists
Histamine.Antagonists
Histamine.H1.Antagonists
Serine.Proteinase.Inhibitors
Adrenergic.alpha.Agonists
Dopamine.Antagonists
Adrenergic.alpha.Antagonists
Cholinergic.Antagonists
Muscarinic.Antagonists
Serotonin.Receptor.Agonists
-0.4 -0.2 0.0 0.2
Location Shift
Percent of Drugs in Class Approved for Indication
0.125
0.100
0.075
0.050
0.025
0.000
Lung Cancer
C
Protein.Kinase.Inhibitors
Kinase.Inhibitor
Topoisomerase.Inhibitors
Topoisomerase.II.Inhibitors
BCRP.ABCG2.Inhibitors
BSEP.ABCB11.Inhibitors
Hormone.Antagonists
Cyclooxygenase.Inhibitors
Hormones..Hormone.Substitutes..and.Hormone.Antagonists
Cytochrome.P.450.CYP2C8.Inhibitors
Cytochrome.P.450.CYP1A2.Inhibitors
Combined.Inhibitors.of.CYP3A4.and.P.glycoprotein
Cytochrome.P.450.CYP2B6.Inhibitors
Cytochrome.P.450.CYP2C9.Inhibitors
Cytochrome.P.450.CYP2A6.Inhibitors
P.glycoprotein.ABCB1.Inhibitors
Enzyme.Inhibitors
Nucleic.Acid.Synthesis.Inhibitors
Cytochrome.P.450.CYP3A.Inhibitors
Cytochrome.P.450.Enzyme.Inhibitors
Cytochrome.P.450.CYP3A4.Inhibitors
Protein.Synthesis.Inhibitors
Adrenergic.alpha.Agonists
Dopamine.Antagonists
Protease.Inhibitors
Monoamine.Oxidase.Inhibitors
Beta2.Agonists
Phosphodiesterase.Inhibitors
Adrenergic.Agonists
Adrenergic.alpha.Antagonists
Neurotransmitter.Uptake.Inhibitors
Adrenergic.Uptake.Inhibitors
Serotonin.Receptor.Agonists
Histamine.H1.Antagonists
Histamine.Antagonists
Adrenergic.beta.Agonists
Adrenergic.Antagonists
Serine.Proteinase.Inhibitors
Adrenergic.beta.Antagonists
Cholinergic.Antagonists
Muscarinic.Antagonists
-0.5 0.0 0.5 1.0
Location Shift
Percent of Drugs in Class Approved for Indication
0.25
0.20
0.15
0.10
0.05
0.00
Breast Cancer
D
Protein.Kinase.Inhibitors
Kinase.Inhibitor
Hormone.Antagonists
Cyclooxygenase.Inhibitors
Topoisomerase.Inhibitors
BCRP.ABCG2.Inhibitors
BSEP.ABCB11.Inhibitors
Topoisomerase.II.Inhibitors
Hormones..Hormone.Substitutes..and.Hormone.Antagonists
Combined.Inhibitors.of.CYP3A4.and.P.glycoprotein
Cytochrome.P.450.CYP2C8.Inhibitors
Cytochrome.P.450.CYP2A6.Inhibitors
Enzyme.Inhibitors
Cytochrome.P.450.Enzyme.Inhibitors
Cytochrome.P.450.CYP2B6.Inhibitors
Cytochrome.P.450.CYP2C9.Inhibitors
P.glycoprotein.ABCB1.Inhibitors
Cytochrome.P.450.CYP3A.Inhibitors
Nucleic.Acid.Synthesis.Inhibitors
Cytochrome.P.450.CYP1A2.Inhibitors
Cytochrome.P.450.CYP3A4.Inhibitors
Cytochrome.P.450.CYP2C19.Inhibitors
Protein.Synthesis.Inhibitors
Cytochrome.P.450.CYP2E1.Inhibitors
Protease.Inhibitors
Cytochrome.P.450.CYP2D6.Inhibitors
Alpha1.Antagonists
Adrenergic.alpha.Antagonists
Platelet.Aggregation.Inhibitors
Neurotransmitter.Uptake.Inhibitors
Adrenergic.alpha.Agonists
Adrenergic.Uptake.Inhibitors
Histamine.H1.Antagonists
Serine.Proteinase.Inhibitors
Phosphodiesterase.Inhibitors
Antiemetics.Antagonists
Histamine.Antagonists
Dopamine.Antagonists
Adrenergic.Agonists
Serotonin.Antagonists
Beta2.Agonists
Adrenergic.Antagonists
Adrenergic.beta.Agonists
Serotonin.Receptor.Agonists
Cholinergic.Antagonists
Adrenergic.beta.Antagonists
Muscarinic.Antagonists
-0.5 0.0 0.5
Location Shift
Percent of Drugs in Class Approved for Indication
0.10
0.05
0.00
Thryoid Cancer
E
Topoisomerase.Inhibitors
Topoisomerase.II.Inhibitors
Serine.Proteinase.Inhibitors
Cytochrome.P.450.CYP2C8.Inhibitors
Adrenergic.beta.Antagonists
Adrenergic.beta.Agonists
Cytochrome.P.450.Enzyme.Inhibitors
Adrenergic.Agonists
Hormones..Hormone.Substitutes..and.Hormone.Antagonists
Adrenergic.Antagonists
BSEP.ABCB11.Inhibitors
Beta2.Agonists
Combined.Inhibitors.of.CYP3A4.and.P.glycoprotein
Phosphodiesterase.Inhibitors
Dopamine.Antagonists
Cytochrome.P.450.CYP1A2.Inhibitors
Adrenergic.alpha.Agonists
Hormone.Antagonists
Serotonin.Antagonists
Cytochrome.P.450.CYP2C9.Inhibitors
Adrenergic.alpha.Antagonists
Enzyme.Inhibitors
Protease.Inhibitors
Histamine.Antagonists
BCRP.ABCG2.Inhibitors
Cyclooxygenase.Inhibitors
Cytochrome.P.450.CYP2C19.Inhibitors
Histamine.H1.Antagonists
Cytochrome.P.450.CYP2A6.Inhibitors
Nucleic.Acid.Synthesis.Inhibitors
Alpha1.Antagonists
Cytochrome.P.450.CYP2B6.Inhibitors
P.glycoprotein.ABCB1.Inhibitors
Cytochrome.P.450.CYP3A.Inhibitors
Cytochrome.P.450.CYP3A4.Inhibitors
Cytochrome.P.450.CYP2D6.Inhibitors
Protein.Synthesis.Inhibitors
Monoamine.Oxidase.Inhibitors
Cholinergic.Antagonists
Muscarinic.Antagonists
-0.2 0.0 0.2 0.4
Location Shift
Percent of Drugs in Class Approved for Indication
0.05
0.04
0.03
0.02
0.01
0.00
Large Intestine Cancer
F
Protein.Kinase.Inhibitors
Kinase.Inhibitor
Topoisomerase.Inhibitors
Topoisomerase.II.Inhibitors
Cyclooxygenase.Inhibitors
BSEP.ABCB11.Inhibitors
BCRP.ABCG2.Inhibitors
Enzyme.Inhibitors
Nucleic.Acid.Synthesis.Inhibitors
Cytochrome.P.450.CYP2C9.Inhibitors
Cytochrome.P.450.CYP2D6.Inhibitors
Adrenergic.alpha.Agonists
Cytochrome.P.450.CYP2B6.Inhibitors
Alpha1.Antagonists
Cytochrome.P.450.CYP2C19.Inhibitors
Cytochrome.P.450.CYP2E1.Inhibitors
Cytochrome.P.450.CYP2A6.Inhibitors
Histamine.H1.Antagonists
Neurotransmitter.Uptake.Inhibitors
Serotonin.Antagonists
Serine.Proteinase.Inhibitors
Cholinesterase.Inhibitors
Antiemetics.Antagonists
Histamine.Antagonists
Dopamine.Antagonists
Adrenergic.Agonists
Cholinergic.Antagonists
Adrenergic.Uptake.Inhibitors
Muscarinic.Antagonists
Serotonin.Receptor.Agonists
Adrenergic.Antagonists
Beta2.Agonists
Adrenergic.beta.Agonists
Adrenergic.beta.Antagonists
-0.50 -0.25 0.00 0.25 0.50
Location Shift
Percent of Drugs in Class Approved for Indication
0.05
0.04
0.03
0.02
0.01
0.00
Upper Aerodigestive tract Cancer
G
Protein.Kinase.Inhibitors
BSEP.ABCB11.Inhibitors
BCRP.ABCG2.Inhibitors
Phosphodiesterase.Inhibitors
Kinase.Inhibitor
Cytochrome.P.450.CYP2C8.Inhibitors
Combined.Inhibitors.of.CYP3A4.and.P.glycoprotein
Topoisomerase.Inhibitors
Cytochrome.P.450.Enzyme.Inhibitors
Cytochrome.P.450.CYP2C9.Inhibitors
Cytochrome.P.450.CYP3A.Inhibitors
Topoisomerase.II.Inhibitors
Cytochrome.P.450.CYP3A4.Inhibitors
Cytochrome.P.450.CYP1A2.Inhibitors
Cytochrome.P.450.CYP2E1.Inhibitors
P.glycoprotein.ABCB1.Inhibitors
Alpha1.Antagonists
Adrenergic.alpha.Antagonists
Protease.Inhibitors
Cytochrome.P.450.CYP2B6.Inhibitors
Cytochrome.P.450.CYP2A6.Inhibitors
Cytochrome.P.450.CYP2D6.Inhibitors
Enzyme.Inhibitors
Cytochrome.P.450.CYP2C19.Inhibitors
Histamine.H1.Antagonists
Protein.Synthesis.Inhibitors
Hormones..Hormone.Substitutes..and.Hormone.Antagonists
Neurotransmitter.Uptake.Inhibitors
Adrenergic.alpha.Agonists
Serine.Proteinase.Inhibitors
Beta2.Agonists
Adrenergic.Agonists
Adrenergic.beta.Antagonists
Adrenergic.beta.Agonists
Cholinergic.Antagonists
Muscarinic.Antagonists
-0.4 0.0 0.4
Location Shift
Percent of Drugs in Class Approved for Indication
0.03
0.02
0.01
0.00
Gastric Cancer
H
Topoisomerase.II.Inhibitors
Protein.Kinase.Inhibitors
Kinase.Inhibitor
Topoisomerase.Inhibitors
Cyclooxygenase.Inhibitors
Cytochrome.P.450.Enzyme.Inhibitors
BSEP.ABCB11.Inhibitors
BCRP.ABCG2.Inhibitors
Hormone.Antagonists
Cytochrome.P.450.CYP2C8.Inhibitors
Alpha2.Agonists
Enzyme.Inhibitors
Cytochrome.P.450.CYP1A2.Inhibitors
Alpha1.Antagonists
Combined.Inhibitors.of.CYP3A4.and.P.glycoprotein
Cytochrome.P.450.CYP2C9.Inhibitors
Cytochrome.P.450.CYP2E1.Inhibitors
Hormones..Hormone.Substitutes..and.Hormone.Antagonists
Cytochrome.P.450.CYP2A6.Inhibitors
Cytochrome.P.450.CYP2B6.Inhibitors
Protease.Inhibitors
Dopamine.Antagonists
Nucleic.Acid.Synthesis.Inhibitors
Cytochrome.P.450.CYP2C19.Inhibitors
Platelet.Aggregation.Inhibitors
Cytochrome.P.450.CYP3A.Inhibitors
Cytochrome.P.450.CYP2D6.Inhibitors
Protein.Synthesis.Inhibitors
Cytochrome.P.450.CYP3A4.Inhibitors
P.glycoprotein.ABCB1.Inhibitors
Monoamine.Oxidase.Inhibitors
Serotonin.Receptor.Agonists
Adrenergic.beta.Agonists
Adrenergic.Antagonists
Cholinergic.Antagonists
Adrenergic.beta.Antagonists
Muscarinic.Antagonists
-0.5 0.0 0.5 1.0
Location Shift
Percent of Drugs in Class Approved for Indication
0.100
0.075
0.050
0.025
0.000
Renal Cancer
I
Protein.Kinase.Inhibitors
Kinase.Inhibitor
Topoisomerase.Inhibitors
Topoisomerase.II.Inhibitors
BCRP.ABCG2.Inhibitors
Cyclooxygenase.Inhibitors
BSEP.ABCB11.Inhibitors
Cytochrome.P.450.CYP2C8.Inhibitors
Combined.Inhibitors.of.CYP3A4.and.P.glycoprotein
Enzyme.Inhibitors
Serine.Proteinase.Inhibitors
Cytochrome.P.450.CYP2C9.Inhibitors
Hormone.Antagonists
Cytochrome.P.450.CYP1A2.Inhibitors
P.glycoprotein.ABCB1.Inhibitors
Hormones..Hormone.Substitutes..and.Hormone.Antagonists
Cytochrome.P.450.CYP2A6.Inhibitors
Cytochrome.P.450.Enzyme.Inhibitors
Cytochrome.P.450.CYP3A.Inhibitors
Cytochrome.P.450.CYP3A4.Inhibitors
Phosphodiesterase.Inhibitors
Cytochrome.P.450.CYP2C19.Inhibitors
Protease.Inhibitors
Cytochrome.P.450.CYP2B6.Inhibitors
Platelet.Aggregation.Inhibitors
Cytochrome.P.450.CYP2D6.Inhibitors
Nucleic.Acid.Synthesis.Inhibitors
Cytochrome.P.450.CYP2E1.Inhibitors
Protein.Synthesis.Inhibitors
Histamine.H1.Antagonists
Histamine.Antagonists
Adrenergic.Antagonists
Neurotransmitter.Uptake.Inhibitors
Serotonin.Antagonists
Serotonin.Receptor.Agonists
Adrenergic.beta.Antagonists
Antiemetics.Antagonists
Adrenergic.Uptake.Inhibitors
Cholinergic.Antagonists
Muscarinic.Antagonists
-0.5 0.0 0.5 1.0
Location Shift
Percent of Drugs in Class Approved for Indication
0.15
0.10
0.05
0.00

Figure S7

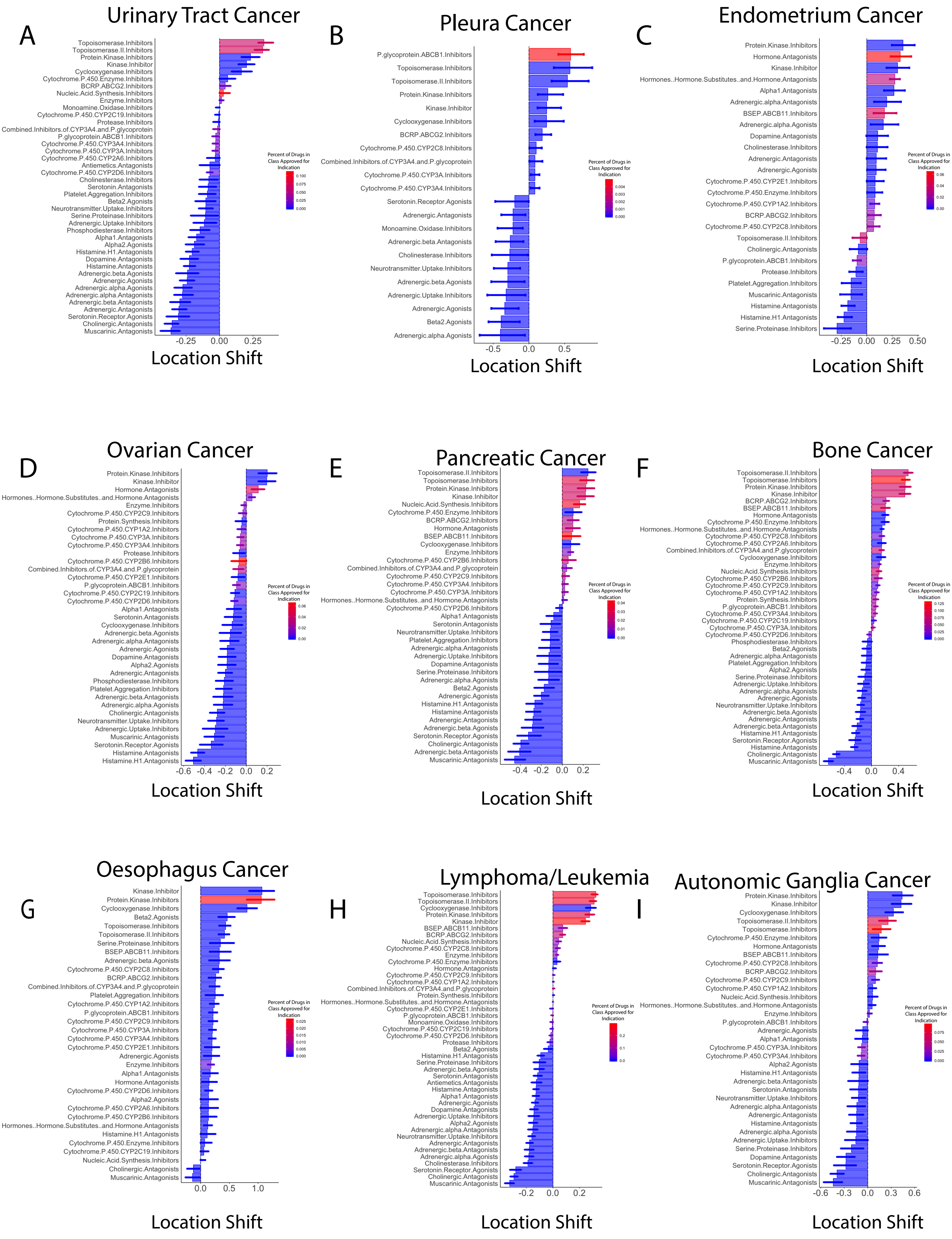
A
Urinary Tract Cancer
Topoisomerase.Inhibitors
Topoisomerase.II.Inhibitors
Protein.Kinase.Inhibitors
Kinase.Inhibitor
Cyclooxygenase.Inhibitors
Cytochrome.P.450.Enzyme.Inhibitors
BCRP.ABCG2.Inhibitors
Nucleic.Acid.Synthesis.Inhibitors
Enzyme.Inhibitors
Monoamine.Oxidase.Inhibitors
Cytochrome.P.450.CYP2C19.Inhibitors
Protease.Inhibitors
Combined.Inhibitors.of.CYP3A4.and.P.glycoprotein
P.glycoprotein.ABCB1.Inhibitors
Cytochrome.P.450.CYP3A4.Inhibitors
Cytochrome.P.450.CYP3A.Inhibitors
Cytochrome.P.450.CYP2A6.Inhibitors
Antiemetics.Antagonists
Cytochrome.P.450.CYP2D6.Inhibitors
Cholinesterase.Inhibitors
Serotonin.Antagonists
Platelet.Aggregation.Inhibitors
Beta2.Agonists
Neurotransmitter.Uptake.Inhibitors
Serine.Proteinase.Inhibitors
Adrenergic.Uptake.Inhibitors
Phosphodiesterase.Inhibitors
Alpha1.Antagonists
Alpha2.Agonists
Histamine.H1.Antagonists
Dopamine.Antagonists
Histamine.Antagonists
Adrenergic.beta.Agonists
Adrenergic.Agonists
Adrenergic.alpha.Agonists
Adrenergic.alpha.Antagonists
Adrenergic.beta.Antagonists
Adrenergic.Antagonists
Serotonin.Receptor.Agonists
Cholinergic.Antagonists
Muscarinic.Antagonists
-0.25
0.00
0.25
Location Shift
Percent of Drugs in Class Approved for Indication
0.100
0.075
0.050
0.025
0.000
B
Pleura Cancer
P.glycoprotein.ABCB1.Inhibitors
Topoisomerase.Inhibitors
Topoisomerase.II.Inhibitors
Protein.Kinase.Inhibitors
Kinase.Inhibitor
Cyclooxygenase.Inhibitors
BCRP.ABCG2.Inhibitors
Cytochrome.P.450.CYP2C8.Inhibitors
Combined.Inhibitors.of.CYP3A4.and.P.glycoprotein
Cytochrome.P.450.CYP3A.Inhibitors
Cytochrome.P.450.CYP3A4.Inhibitors
Serotonin.Receptor.Agonists
Adrenergic.Antagonists
Monoamine.Oxidase.Inhibitors
Adrenergic.beta.Antagonists
Cholinesterase.Inhibitors
Neurotransmitter.Uptake.Inhibitors
Adrenergic.beta.Agonists
Adrenergic.Uptake.Inhibitors
Adrenergic.Agonists
Beta2.Agonists
Adrenergic.alpha.Agonists
-0.5
0.0
0.5
Location Shift
Percent of Drugs in Class Approved for Indication
0.004
0.003
0.002
0.001
0.000
C
Endometrium Cancer
Protein.Kinase.Inhibitors
Hormone.Antagonists
Kinase.Inhibitor
Hormones..Hormone.Substitutes..and.Hormone.Antagonists
Alpha1.Antagonists
Adrenergic.alpha.Antagonists
BSEP.ABCB11.Inhibitors
Adrenergic.alpha.Agonists
Dopamine.Antagonists
Cholinesterase.Inhibitors
Adrenergic.Antagonists
Adrenergic.Agonists
Cytochrome.P.450.CYP2E1.Inhibitors
Cytochrome.P.450.Enzyme.Inhibitors
Cytochrome.P.450.CYP1A2.Inhibitors
BCRP.ABCG2.Inhibitors
Cytochrome.P.450.CYP2C8.Inhibitors
Topoisomerase.II.Inhibitors
Cholinergic.Antagonists
P.glycoprotein.ABCB1.Inhibitors
Protease.Inhibitors
Platelet.Aggregation.Inhibitors
Muscarinic.Antagonists
Histamine.Antagonists
Histamine.H1.Antagonists
Serine.Proteinase.Inhibitors
-0.25
0.00
0.25
0.50
Location Shift
Percent of Drugs in Class Approved for Indication
0.06
0.04
0.02
0.00
D
Ovarian Cancer
Protein.Kinase.Inhibitors
Kinase.Inhibitor
Hormone.Antagonists
Hormones..Hormone.Substitutes..and.Hormone.Antagonists
Enzyme.Inhibitors
Cytochrome.P.450.CYP2C9.Inhibitors
Protein.Synthesis.Inhibitors
Cytochrome.P.450.CYP1A2.Inhibitors
Cytochrome.P.450.CYP3A.Inhibitors
Cytochrome.P.450.CYP3A4.Inhibitors
Protease.Inhibitors
Cytochrome.P.450.CYP2B6.Inhibitors
Combined.Inhibitors.of.CYP3A4.and.P.glycoprotein
Cytochrome.P.450.CYP2E1.Inhibitors
P.glycoprotein.ABCB1.Inhibitors
Cytochrome.P.450.CYP2C19.Inhibitors
Cytochrome.P.450.CYP2D6.Inhibitors
Alpha1.Antagonists
Serotonin.Antagonists
Cyclooxygenase.Inhibitors
Adrenergic.beta.Agonists
Adrenergic.alpha.Antagonists
Adrenergic.Agonists
Dopamine.Antagonists
Alpha2.Agonists
Adrenergic.Antagonists
Phosphodiesterase.Inhibitors
Platelet.Aggregation.Inhibitors
Adrenergic.beta.Antagonists
Adrenergic.alpha.Agonists
Cholinergic.Antagonists
Neurotransmitter.Uptake.Inhibitors
Adrenergic.Uptake.Inhibitors
Muscarinic.Antagonists
Serotonin.Receptor.Agonists
Histamine.Antagonists
Histamine.H1.Antagonists
-0.6
-0.4
-0.2
0.0
0.2
Location Shift
Percent of Drugs in Class Approved for Indication
0.06
0.04
0.02
0.00
E
Pancreatic Cancer
Topoisomerase.II.Inhibitors
Topoisomerase.Inhibitors
Protein.Kinase.Inhibitors
Kinase.Inhibitor
Nucleic.Acid.Synthesis.Inhibitors
Cytochrome.P.450.Enzyme.Inhibitors
BCRP.ABCG2.Inhibitors
Hormone.Antagonists
BSEP.ABCB11.Inhibitors
Cyclooxygenase.Inhibitors
Enzyme.Inhibitors
Cytochrome.P.450.CYP2B6.Inhibitors
Combined.Inhibitors.of.CYP3A4.and.P.glycoprotein
Cytochrome.P.450.CYP2C9.Inhibitors
Cytochrome.P.450.CYP3A4.Inhibitors
Cytochrome.P.450.CYP3A.Inhibitors
Hormones..Hormone.Substitutes..and.Hormone.Antagonists
Cytochrome.P.450.CYP2D6.Inhibitors
Alpha1.Antagonists
Serotonin.Antagonists
Neurotransmitter.Uptake.Inhibitors
Platelet.Aggregation.Inhibitors
Adrenergic.alpha.Antagonists
Adrenergic.Uptake.Inhibitors
Dopamine.Antagonists
Serine.Proteinase.Inhibitors
Adrenergic.alpha.Agonists
Beta2.Agonists
Adrenergic.Agonists
Histamine.H1.Antagonists
Histamine.Antagonists
Adrenergic.Antagonists
Adrenergic.beta.Agonists
Serotonin.Receptor.Agonists
Cholinergic.Antagonists
Adrenergic.beta.Antagonists
Muscarinic.Antagonists
-0.4
-0.2
0.0
0.2
Location Shift
Percent of Drugs in Class Approved for Indication
0.04
0.03
0.02
0.01
0.00
F
Bone Cancer
Topoisomerase.II.Inhibitors
Topoisomerase.Inhibitors
Protein.Kinase.Inhibitors
Kinase.Inhibitor
BCRP.ABCG2.Inhibitors
BSEP.ABCB11.Inhibitors
Hormone.Antagonists
Cytochrome.P.450.Enzyme.Inhibitors
Hormones..Hormone.Substitutes..and.Hormone.Antagonists
Cytochrome.P.450.CYP2C8.Inhibitors
Cytochrome.P.450.CYP2A6.Inhibitors
Combined.Inhibitors.of.CYP3A4.and.P.glycoprotein
Cyclooxygenase.Inhibitors
Enzyme.Inhibitors
Nucleic.Acid.Synthesis.Inhibitors
Cytochrome.P.450.CYP2B6.Inhibitors
Cytochrome.P.450.CYP2C9.Inhibitors
Cytochrome.P.450.CYP1A2.Inhibitors
Protein.Synthesis.Inhibitors
P.glycoprotein.ABCB1.Inhibitors
Cytochrome.P.450.CYP3A4.Inhibitors
Cytochrome.P.450.CYP2C19.Inhibitors
Cytochrome.P.450.CYP3A.Inhibitors
Cytochrome.P.450.CYP2D6.Inhibitors
Phosphodiesterase.Inhibitors
Beta2.Agonists
Adrenergic.alpha.Antagonists
Platelet.Aggregation.Inhibitors
Alpha2.Agonists
Serine.Proteinase.Inhibitors
Adrenergic.Uptake.Inhibitors
Adrenergic.alpha.Agonists
Adrenergic.Agonists
Neurotransmitter.Uptake.Inhibitors
Adrenergic.beta.Agonists
Adrenergic.Antagonists
Adrenergic.beta.Antagonists
Histamine.H1.Antagonists
Serotonin.Receptor.Agonists
Histamine.Antagonists
Cholinergic.Antagonists
Muscarinic.Antagonists
-0.4
0.0
0.4
Location Shift
Percent of Drugs in Class Approved for Indication
0.125
0.100
0.075
0.050
0.025
0.000
G
Oesophagus Cancer
Kinase.Inhibitor
Protein.Kinase.Inhibitors
Cyclooxygenase.Inhibitors
Beta2.Agonists
Topoisomerase.Inhibitors
Topoisomerase.II.Inhibitors
Serine.Proteinase.Inhibitors
BSEP.ABCB11.Inhibitors
Adrenergic.beta.Agonists
Cytochrome.P.450.CYP2C8.Inhibitors
BCRP.ABCG2.Inhibitors
Combined.Inhibitors.of.CYP3A4.and.P.glycoprotein
Platelet.Aggregation.Inhibitors
Cytochrome.P.450.CYP1A2.Inhibitors
P.glycoprotein.ABCB1.Inhibitors
Cytochrome.P.450.CYP2C9.Inhibitors
Cytochrome.P.450.CYP3A.Inhibitors
Cytochrome.P.450.CYP3A4.Inhibitors
Cytochrome.P.450.CYP2E1.Inhibitors
Adrenergic.Agonists
Enzyme.Inhibitors
Alpha1.Antagonists
Hormone.Antagonists
Cytochrome.P.450.CYP2D6.Inhibitors
Alpha2.Agonists
Cytochrome.P.450.CYP2A6.Inhibitors
Cytochrome.P.450.CYP2B6.Inhibitors
Hormones..Hormone.Substitutes..and.Hormone.Antagonists
Histamine.H1.Antagonists
Cytochrome.P.450.Enzyme.Inhibitors
Cytochrome.P.450.CYP2C19.Inhibitors
Nucleic.Acid.Synthesis.Inhibitors
Cholinergic.Antagonists
Muscarinic.Antagonists
0.0
0.5
1.0
Location Shift
Percent of Drugs in Class Approved for Indication
0.025
0.020
0.015
0.010
0.005
0.000
H
Lymphoma/Leukemia
Topoisomerase.Inhibitors
Topoisomerase.II.Inhibitors
Cyclooxygenase.Inhibitors
Protein.Kinase.Inhibitors
Kinase.Inhibitor
BSEP.ABCB11.Inhibitors
BCRP.ABCG2.Inhibitors
Nucleic.Acid.Synthesis.Inhibitors
Cytochrome.P.450.CYP2C8.Inhibitors
Enzyme.Inhibitors
Cytochrome.P.450.Enzyme.Inhibitors
Hormone.Antagonists
Cytochrome.P.450.CYP2C9.Inhibitors
Cytochrome.P.450.CYP1A2.Inhibitors
Combined.Inhibitors.of.CYP3A4.and.P.glycoprotein
Protein.Synthesis.Inhibitors
Hormones..Hormone.Substitutes..and.Hormone.Antagonists
Cytochrome.P.450.CYP2E1.Inhibitors
P.glycoprotein.ABCB1.Inhibitors
Monoamine.Oxidase.Inhibitors
Cytochrome.P.450.CYP2C19.Inhibitors
Cytochrome.P.450.CYP2D6.Inhibitors
Protease.Inhibitors
Beta2.Agonists
Histamine.H1.Antagonists
Serine.Proteinase.Inhibitors
Adrenergic.beta.Agonists
Serotonin.Antagonists
Antiemetics.Antagonists
Histamine.Antagonists
Alpha1.Antagonists
Adrenergic.Agonists
Dopamine.Antagonists
Adrenergic.Uptake.Inhibitors
Alpha2.Agonists
Adrenergic.alpha.Antagonists
Neurotransmitter.Uptake.Inhibitors
Adrenergic.Antagonists
Adrenergic.beta.Antagonists
Adrenergic.alpha.Agonists
Cholinesterase.Inhibitors
Serotonin.Receptor.Agonists
Cholinergic.Antagonists
Muscarinic.Antagonists
-0.2
0.0
0.2
Location Shift
Percent of Drugs in Class Approved for Indication
0.2
0.1
0.0
I
Autonomic Ganglia Cancer
Protein.Kinase.Inhibitors
Kinase.Inhibitor
Cyclooxygenase.Inhibitors
Topoisomerase.II.Inhibitors
Topoisomerase.Inhibitors
Cytochrome.P.450.Enzyme.Inhibitors
Hormone.Antagonists
BSEP.ABCB11.Inhibitors
Cytochrome.P.450.CYP2C8.Inhibitors
BCRP.ABCG2.Inhibitors
Cytochrome.P.450.CYP2C9.Inhibitors
Cytochrome.P.450.CYP1A2.Inhibitors
Nucleic.Acid.Synthesis.Inhibitors
Hormones..Hormone.Substitutes..and.Hormone.Antagonists
Enzyme.Inhibitors
P.glycoprotein.ABCB1.Inhibitors
Adrenergic.Agonists
Alpha1.Antagonists
Cytochrome.P.450.CYP3A.Inhibitors
Cytochrome.P.450.CYP3A4.Inhibitors
Alpha2.Agonists
Histamine.H1.Antagonists
Adrenergic.beta.Antagonists
Serotonin.Antagonists
Neurotransmitter.Uptake.Inhibitors
Adrenergic.alpha.Antagonists
Adrenergic.Antagonists
Histamine.Antagonists
Adrenergic.alpha.Agonists
Adrenergic.Uptake.Inhibitors
Serine.Proteinase.Inhibitors
Dopamine.Antagonists
Serotonin.Receptor.Agonists
Cholinergic.Antagonists
Muscarinic.Antagonists
-0.6
-0.3
0.0
0.3
0.6
Location Shift
Percent of Drugs in Class Approved for Indication
0.075
0.050
0.025
0.000

Figure S8

A

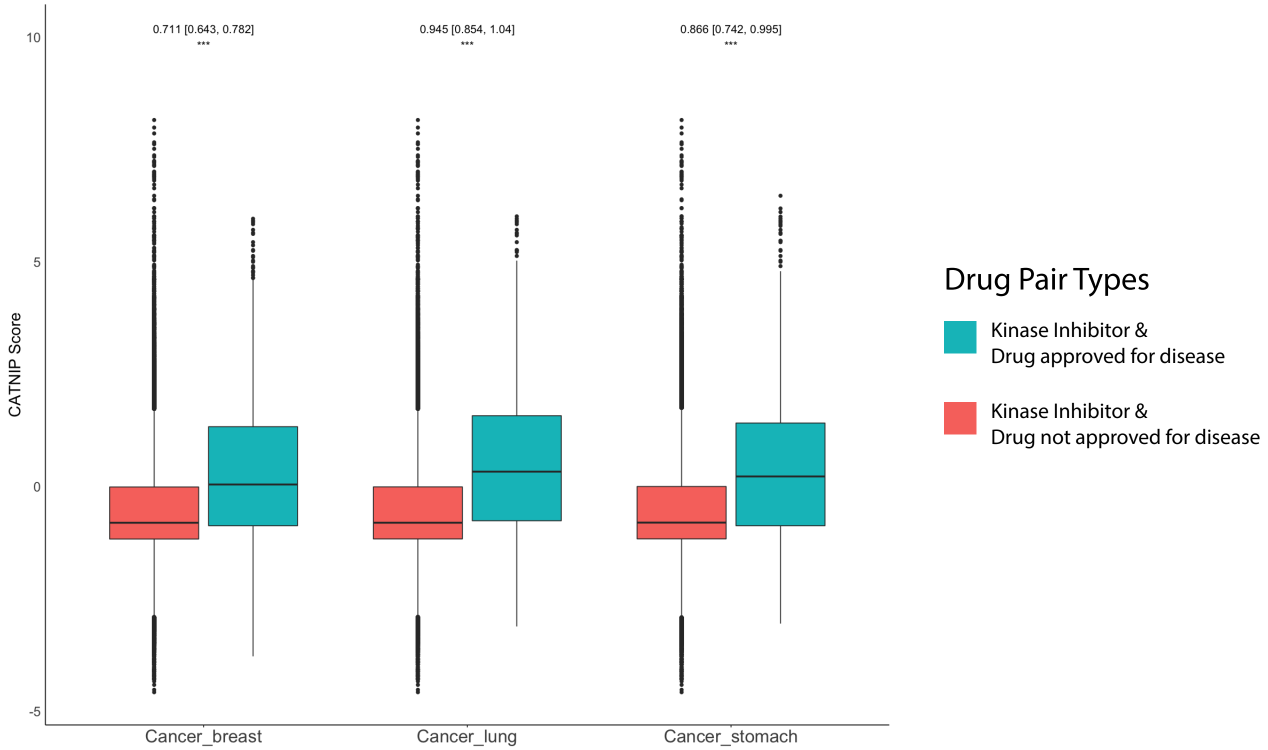
0.711 [0.643, 0.782]
***
0.945 [0.854, 1.04]
***
0.866 [0.742, 0.995]
***
10
5
0
-5
CATNIP Score
Cancer_breast
Cancer_lung
Cancer_stomach
Drug Pair Types
Kinase Inhibitor & Drug approved for disease
Kinase Inhibitor & Drug not approved for disease

A

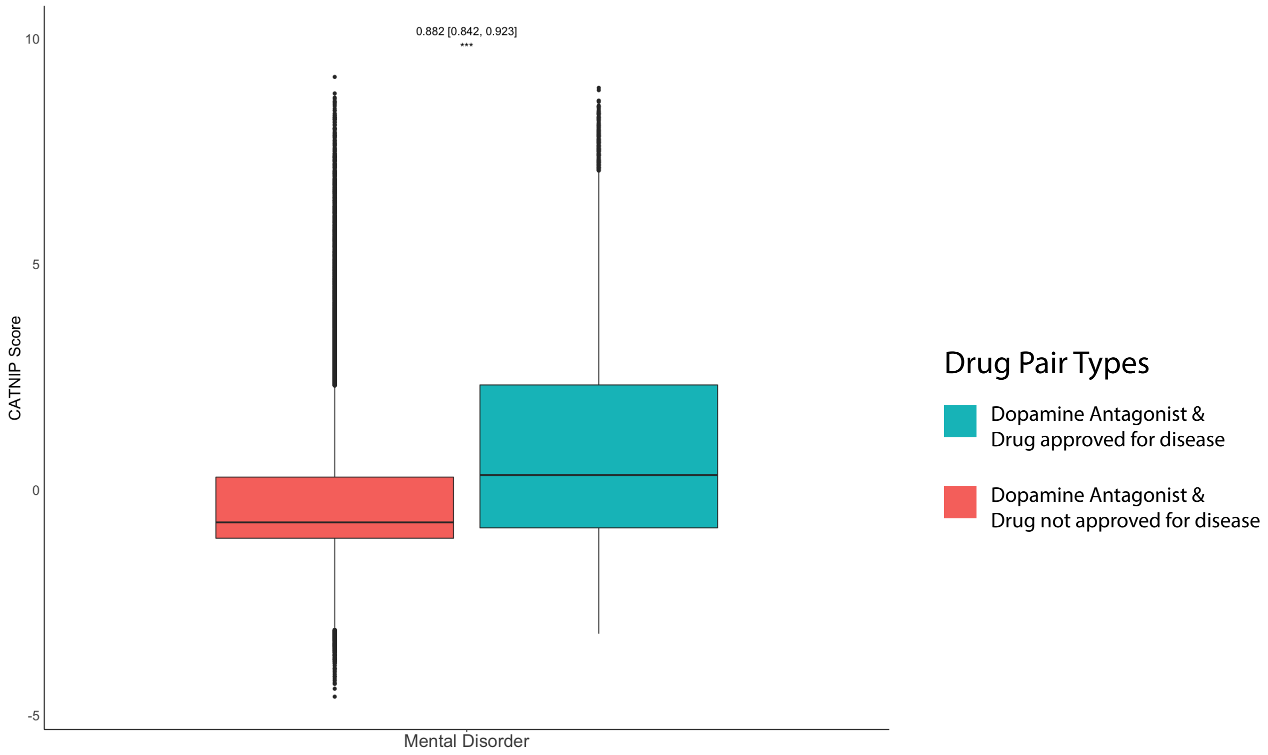
0.882 [0.842, 0.923]
***
10
5
0
-5
CATNIP Score
Mental Disorder
Drug Pair Types
Dopamine Antagonist & Drug approved for disease
Dopamine Antagonist & Drug not approved for disease

Figure S9

A

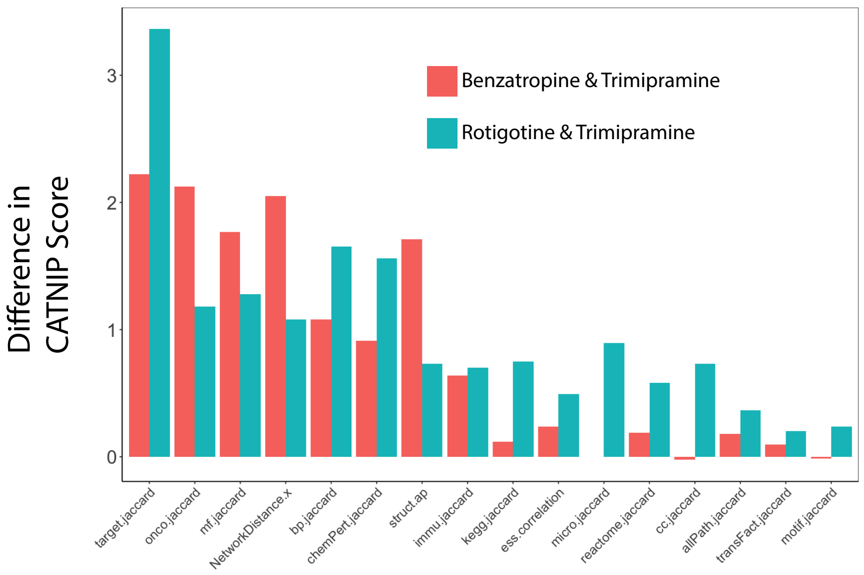

Benzatropine & Trimipramine
Rotigotine & Trimipramine
Difference in CATNIP Score
0
1
2
3
target.jaccard
onco.jaccard
mf.jaccard
NetworkDistance.x
bp.jaccard
chemPert.jaccard
struct.ap
immu.jaccard
kegg.jaccard
ess.correlation
micro.jaccard
reactome.jaccard
cc.jaccard
allPath.jaccard
transFact.jaccard
motif.jaccard


Figure S10

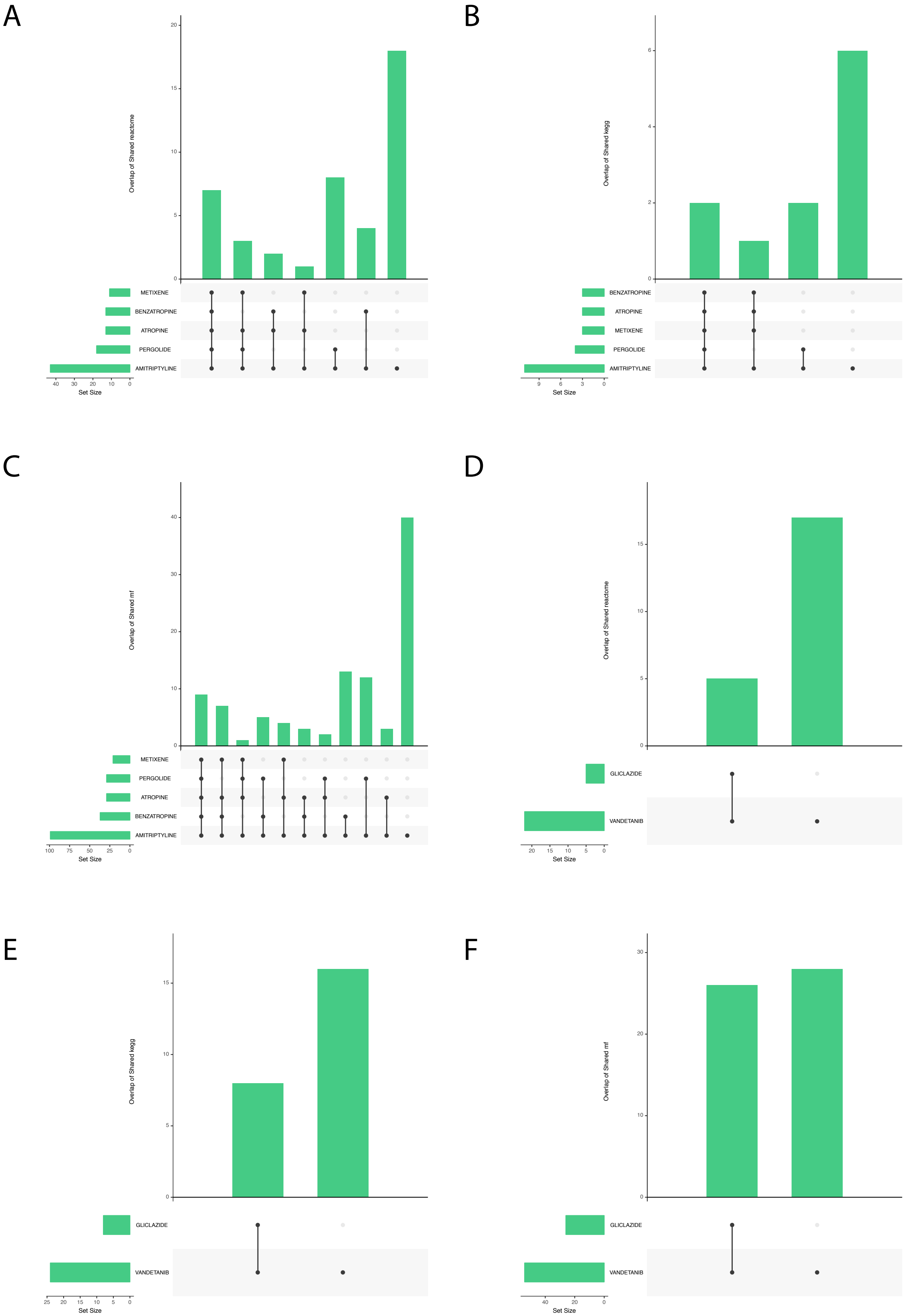
A
Overlap of Shared reactome
0
5
10
15
20
METIXENE
BENZATROPINE
ATROPINE
PERGOLIDE
AMITRIPTYLINE
40 30 20 10 0
Set Size
B
Overlap of Shared kegg
0
2
4
6
BENZATROPINE
ATROPINE
METIXENE
PERGOLIDE
AMITRIPTYLINE
9 6 3 0
Set Size
C
Overlap of Shared mf
0
10
20
30
40
METIXENE
PERGOLIDE
ATROPINE
BENZATROPINE
AMITRIPTYLINE
100 75 50 25 0
Set Size
D
Overlap of Shared reactome
0
5
10
15
GLICLAZIDE
VANDETANIB
20 15 10 5 0
Set Size
E
Overlap of Shared kegg
0
5
10
15
GLICLAZIDE
VANDETANIB
25 20 15 10 5 0
Set Size
F
Overlap of Shared mf
0
10
20
30
GLICLAZIDE
VANDETANIB
40 20 0
Set Size

Figure S10

| Drug Property | Similarity Metric | Source |
|---|---|---|
| Drug Molecular Target | Jaccard Index | DrugBank |
| Protein-Protein Network | Target Distance | In-house PPI Network (**Methods**) |
| GO: Biological Process | Jaccard Index | MSigDB |
| GO: Molecular Function | Jaccard Index | MSigDB |
| GO: Cellular Component | Jaccard Index | MSigDB |
| Transcription Factor | Jaccard Index | MSigDB |
| KEGG Pathways | Jaccard Index | MSigDB |
| Reactome Pathways | Jaccard Index | MSigDB |
| Target Essentiality | Jaccard Index | MSigDB |
| Motif | Jaccard Index | MSigDB |
| Canonical Pathways | Jaccard Index | MSigDB |
| microRNA | Jaccard Index | MSigDB |
| Oncogenic Signature | Jaccard Index | MSigDB |
| Immunogenic Signature | Jaccard Index | MSigDB |
| Chemical Perturbation | Jaccard Index | MSigDB |
| Chemical Structure | Dice Fingerprint Similarity | DrugBank |

Table S1

| | AUC | AUPRC |
|---|---|---|
| **SVM - RADIAL KERNEL** | 0.696 | 0.0557 |
| **LOGISTIC REGRESSION – ELASTIC NET** | 0.662 | 0.0842 |
| **LOGISTIC REGRESSION – LASSO** | 0.662 | 0.0836 |
| **GRADIENT BOOSTING** | 0.841 | 0.189 |

Table S2
